# Quantification of neuron types in the rodent hippocampal formation by data mining and numerical optimization

**DOI:** 10.1101/2021.09.20.460986

**Authors:** Sarojini M. Attili, Keivan Moradi, Diek W. Wheeler, Giorgio A. Ascoli

## Abstract

Quantifying the population sizes of distinct neuron types in different anatomical regions is an essential step towards establishing a brain cell census. Although estimates exist for the total neuronal populations in different species, the number and definition of each specific neuron type are still intensively investigated. Hippocampome.org is an open-source knowledge base with morphological, physiological, and molecular information for 122 neuron types in the rodent hippocampal formation. While such framework identifies all known neuron types in this system, their relative abundances remain largely unknown. This work quantitatively estimates the counts of all Hippocampome.org neuron types by literature mining and numerical optimization. We report the number of neurons in each type identified by main neurotransmitter (glutamate or GABA) and axonal-dendritic patterns throughout 26 subregions and layers of the dentate gyrus, Ammon’s horn, subiculum, and entorhinal cortex. We produce by sensitivity analysis reliable numerical ranges for each type and summarize the amounts across broad neuronal families defined by biomarkers expression and firing dynamics. Study of density distributions indicates that the number of dendritic-targeting interneurons, but not of other neuronal classes, is independent of anatomical volumes. All extracted values, experimental evidence, and related software code are released on Hippocampome.org.

## Introduction

The brain is a highly complex organ encompassing an extraordinary quantity and diversity of cells. Compared to the high-level functional characterization of brain anatomy (Taubert et al., 2010; Lange et al., 1997), knowledge on low-level components such as neuron types remains incomplete. In particular, despite recent progress in single-cell transcriptomics, epigenomic profiling, and microscopic imaging, we still lack a complete census of brain cell types (Mukamel and Ngai, 2019). The US Brain Research through Advancing Innovative Neurotechnologies (BRAIN) Initiative emphasizes mathematical modeling, statistical analysis, and exploratory data mining to accelerate discoveries in this regard (Mott et al., 2018). Building upon this vision, the National Institutes of Health launched the BRAIN Initiative Cell Census Network (BICCN), a consortium of research projects tasked with generating a molecular and anatomical “parts list” at the cellular level within a three-dimensional reference atlas of the mouse brain (Ecker et al., 2017).

The online knowledge repository Hippocampome.org classifies all known neuron types in the hippocampal formation: dentate gyrus (DG), CA3, CA2, CA1, subiculum (Sub), and entorhinal cortex (EC). This resource defines 122 neuron types based on their main neurotransmitter (glutamate or GABA) and the spatial distributions of their axons and dendrites (Wheeler et al., 2015). Hippocampome.org further annotates every neuron type with its reported connectivity (Rees et al., 2016), electrophysiological (Komendantov et al., 2019), molecular (White et al., 2020), synaptic (Moradi et al., 2020), morphological (Tecuatl et al., 2021), and functional (Sanchez-Aguilera et al., 2021) properties, in all cases providing links to the underlying experimental evidence. The ultimate goal of Hippocampome.org is to create biologically plausible computational models of the hippocampus (Venkadesh et al., 2019). Towards this aim, one key piece of information needed is the count of neurons in each type. The present study derives and publicly releases the first estimates of these values using numerical optimization of data mined from the existing scientific literature.

More extensive research is typically available on principal cells than on interneurons (Cembrowski et al., 2016; Faber et al., 2001; Canto and Witter, 2012; Goebbels et al., 2006; Tahvildari and Alonso, 2005; Suzuki and Bekkers, 2011; Ehrlich et al., 2012). Among the hippocampal subregions, DG and CA1 have been most widely researched (Buckmaster et al., 1992; Han et al., 1994; Lubke et al., 1998; Mott et al., 1997; Ramsden et al., 2003; Vida et al., 1998; Svoboda et al., 1999) followed by CA3 and EC (Rapp and Gallagher, 1996; Kaae et al., 2012; Lister et al., 2006; Losonczy et al., 2004; Szabadics et al., 2010; Mercer et al., 2007). Sub and CA2 are the least researched areas of the hippocampal formation (Harris and Stewart, 2000; Mulders et al., 1997; Andrade 2000; Bjerke et al., 2021).

Traditional methods of quantification, such as unbiased stereology, are generally used to report layer-based total counts (Rasmussen et al., 1996; Murakami et al., 2018; Attili et al., 2019). Selected interneuron counts have also been estimated in computational models (Bezaire et al., 2016). However, many electrophysiological studies describe sampling and post-hoc identification of intracellularly recorded neurons, which can be used to derive ratios between the quantities of individual cell types (McBain et al., 1994; Sik et al., 1994; Buhl et al., 1994; Hajos and Mody 1997; Kohus et al., 2016; Szabo et al., 2014). Other important sources of data are reports of proportions of neurons from specific hippocampal layers that express certain molecules (Kosaka et al., 1987; Kim et al., 2017). Here we transform every such relevant piece of evidence into mathematical relations. We then numerically optimize the resultant set of equations to obtain the population estimates for all neuron types in Hippocampome.org.

## Methods

On a high level, the methodology to derive the population estimates of the 122 neuron types classified by Hippocampome.org consists of three steps: literature mining to identify relevant experimental evidence; equation generation, entailing the transformation of the extracted data; and numerical optimization, which produces the estimated counts for each neuron type of interest.

The literature mining process began with a structured search for relevant sources. For every neuron type, Hippocampome.org offers a core list of scientific sources of information providing supporting evidence for its reported properties. We reviewed each of these articles as well as all their cited references. Moreover, we searched Google Scholar for any publication that cited those core sources. From the union of these ‘cited search’ and ‘citing search’, hundreds of journal articles were mined to identify any references to one or more neuron types defined by Hippocampome.org. As there is no standardized nomenclature of hippocampal neurons, it is common for independent researchers to use different names for the same type or the same name for different types (Hamilton et al., 2016). Hence, the first step of the literature mining phase was to identify the properties of the neurons described in each article and map them to the appropriate type(s) from Hippocampome.org, using the putative neurotransmitter and the locations of the soma, axon, and dendrites.

The rare instances with ambiguous or incomplete morphological patterns were resolved by including molecular expression as additional factors in the identification and mapping process. In all cases, data were limited to rats or mice at least 2-week-old.

In order to translate research data into a computable format, we assigned a variable for each neuron type with exclusive somatic location in every subregion of the hippocampus. In other words, one neuron type corresponds to multiple variables if its soma can be found in more than one layer. For example, per Hippocampome.org, CA1 Basket cells may have their soma in stratum pyramidale (SP) or stratum oriens (SO). In this case, we assigned two distinct variables to this neuron type: one for SP-Basket and for SO-Basket. The reason for this choice is that those variables typically take part in different equations (for example, those representing the count of somata in each layer). Through this process, the 122 neuron types gave rise to 198 variables based on their somatic location. Moreover, in order to account for information regarding neuronal groups in certain layers that did not correspond to any Hippocampome.org neuron type, we added 9 ancillary variables: CA2 stratum lacunosum-moleculare (SLM) excitatory, CA2 SLM inhibitory, CA2 SO inhibitory, CA2 stratum radiatum (SR) inhibitory, CA3 SLM excitatory, Sub polymorphic layer inhibitory, EC deep layer inhibitory, medial EC layer I inhibitory, and lateral EC layer I inhibitory. The assignment of multiple variables per neuron types by somatic layer as described earlier and the addition of 9 ancillary neuronal groups resulted into 207 variables for numerical optimization as follows: 22 in DG, 37 in CA3, 9 in CA2, 64 in CA1, 5 in Sub, and 70 in EC (Table 1). The complete list of variables assigned for neuron types based on this classification system can be found in the Supplementary Material.

**Table 1:** The number of neuron types, variables (including the 9 neuronal groups added), equations by type and the residual errors for the six subregions of the hippocampal formation.

| Subregion | Number of neuron types | Number of variables | Total Number of equations |  |  | Residual Error |
| --- | --- | --- | --- | --- | --- | --- |
|  |  |  | Sum | Ratio | Inequality |  |
| DG | 18 | 22 | 115 |  |  | 12.66% |
|  |  |  | 37 | 28 | 50 |  |
| CA3/CA2 | 30 | 46 | 167 |  |  | 18.65% |
|  |  |  | 38 | 31 | 98 |  |
| CA1 | 40 | 64 | 210 |  |  | 22.03% |
|  |  |  | 43 | 77 | 90 |  |
| SUB | 3 | 5 | 30 |  |  | 11.45% |
|  |  |  | 18 | 1 | 11 |  |
| EC | 31 | 70 | 129 |  |  | 6.22% |
|  |  |  | 50 | 39 | 40 |  |

Data mined from various sources were recorded in a structured format that included the bibliographic reference, location of the excerpt of interest, animal species, and interpretation of the content. Each interpretation was converted into an algebraic equation and normalized to unity for use as part of the objective function. A multiplicative scaling factor of 2.44 was applied to convert mouse data to rat (Herculano-Houzel et al., 2006). Numerical densities were converted into neuronal numbers by multiplying with volumes of the corresponding anatomical parcels (Ero et al., 2018; Tecuatl et al., 2021).

Data sources containing neuron type information mined for this work can be divided in three categories that result in distinct types of equations: stereological sums, electrophysiological ratios, and marker inequalities (Table 2). Unbiased stereology measurements and whole-brain high-throughput image-processing methods yielded quantified estimates of neuron counts, where the reported numbers represent totals in a given anatomical parcel (Grady et al., 2003), such as a particular layer of a hippocampal subregion. For example, a study utilizing the optical fractionator reported the presence of 324,000 cells in CA1 pyramidal layer (Hosseini-Sharifabad and Nyengaard, 2007). In these cases, we formulate a linear equation setting the sum of all Hippocampome.org neuron types with soma in CA1 SP to match the reported numerical value.

**Table 2:** Three types of equations obtained from the data transformation process.

| Equation Type | Sum | Ratio | Inequality |
| --- | --- | --- | --- |
| Source | Hosseini-Sharifabad and Nyengaard, 2007 | Gulyas et al., 2010 | Jinno and Kosaka, 2006 |
| Region | CA1 | CA3 | Dentate Gyrus |
| Species | Rat | Mouse | Mouse |
| Evidence excerpt | Table 3 row #6, column 4<br>“Total number of neurons in the granular and pyramidal layers of the rat hippocampus (unilateral values in millions)” | From [...] 61 PV-EGFP cells recorded [...], in 30 cases the [...] boutons were localized to st. pyramidale and only rarely approached ankyrin G-stained profiles (Fig. 1c1–3) suggesting their FSBC origin. Conversely, in the remaining 31 cases the axonal arbor was densest in st. pyramidale and neighboring st. oriens [...] and the boutons formed close appositions with ankyrin G-immunoreactive segments [...], similar to the axon terminals of AAC. | Table 4 row#15, columns 1 & 2<br>“Numerical densities ($\times 10^3 \text{ mm}^{-3}$) of chemically defined GABAergic neuron subtypes in the mouse hippocampus” |
| Location in publication | Page 6, left center | Page 4, left center | Page 12, Left center |
| Interpretation | Total number of neurons in the pyramidal layer of CA1 is 324,000 | Ratio between PV+ basket and axoaxonic cells is 30:31 | Number of parvalbumin positive neurons in granule layer is 3,680 (refer to ‘DG-Biomarkers’ row #6 of supplementary table for data transformation details) |
| Equation | $x_{69} + x_{72} + x_{75} + x_{79} + x_{81} + x_{84} + x_{86} + x_{87} + x_{91} + x_{94} + x_{96} + x_{97} + x_{100} + x_{114} + x_{131} = 324000$ | $\frac{x_6}{x_8} = \frac{31}{30}$ | $x_7 + x_8 \leq 3680$<br>$x_7 + x_8 + x_{17} + x_{21} + x_{22} \geq 3680$ |
| Normalized form | | $\frac{30x_6}{31x_8} - 1 = 0$ | $\frac{x_7 + x_8}{3680} - 1 \leq 0$<br>$\frac{x_7 + x_8 + x_{17} + x_{21} + x_{22}}{3680} - 1 > 0$ |
| | $\begin{aligned} &(x_{69} + x_{72} + x_{75} + x_{79} + x_{81} \\ &\quad + x_{84} + x_{86} + x_{87} \\ &\quad + x_{91} + x_{94} + x_{96} \\ &\quad + x_{97} + x_{100} \\ &\quad + x_{114} \\ &\quad + x_{131})/324000 \\ &- 1 = 0 \end{aligned}$ | | |
| **Least squares form** | $\begin{aligned} &((x_{69} + x_{72} + x_{75} + x_{79} + x_{81} \\ &\quad + x_{84} + x_{86} + x_{87} \\ &\quad + x_{91} + x_{94} + x_{96} \\ &\quad + x_{97} + x_{100} \\ &\quad + x_{114} \\ &\quad + x_{131})/324000 \\ &- 1)^2 = 0 \end{aligned}$ | $\left(\frac{30x_6}{31x_8} - 1\right)^2 = 0$ | $\begin{aligned} &\max\left(0, \frac{x_7 + x_8}{3680} - 1\right)^2 = 0 \\ &\max\left(0, 1 - \frac{x_7 + x_8 + x_{17} + x_{21} + x_{22}}{3680}\right)^2 = 0 \end{aligned}$ |
| **With weights** | $\begin{aligned} &r = 10 \times ((x_{69} + x_{72} + x_{75} + x_{79} \\ &\quad + x_{81} + x_{84} + x_{86} \\ &\quad + x_{87} + x_{91} + x_{94} \\ &\quad + x_{96} + x_{97} \\ &\quad + x_{100} + x_{114} \\ &\quad + x_{131})/324000 \\ &- 1)^2 \end{aligned}$ | $r = 1 \left(\frac{30x_6}{31x_8} - 1\right)^2$ | $\begin{aligned} &r = 1 \times \max\left(0, \frac{x_7 + x_8}{3680} - 1\right)^2 \\ &r = 1 \times \max\left(0, 1 - (x_7 + x_8 + x_{17} \right. \\ &\quad \left. + x_{21} \right. \\ &\quad \left. + x_{22})/3680\right)^2 \end{aligned}$ |

Another source of relevant information comes from numerical reports of intracellularly characterized cells that are morphologically identified post-hoc. From this identification, we derive possible proportions of the corresponding neuron types in that recording location. For example, an electrophysiological study in CA3 SP found that, out of 61 parvalbumin-expressing cells, 30 had axons confined in SP but not targeting the axonal initial segment, whereas 31 had axons concentrated at the SP/SO border and targeting the axonal initial segment (Gulyas et al., 2010). These profiles (and the accompanying illustrations) unambiguously match the Hippocampome.org descriptions of fast-spiking CA3 basket and axo-axonic cells, respectively. Hence, we can formulate an equation setting the ratio of Basket to Axo-axonic cells to 30:31 in CA3. We mined several such electrophysiology-based experimental sources for each subregion.

Several articles published counts or densities of neurons expressing specific molecular biomarkers. We used these numbers to construct inequalities in conjunction with transcriptomics data on Hippocampome.org. Hippocampome.org indicates every neuron type as ‘positive,’ ‘negative,’ or ‘undetermined’ for nearly 100 biomarkers (White et al., 2020). The undetermined cases are largely due to unknown expression of specific neuron types for particular molecules but also include instances of conflicting expression results, location-dependent gradients, and weak signals. To account for undetermined expression, indicating neuron types that might be positive or negative for a given biomarker, we formulated these data into inequalities. For example, the number of parvalbumin (PV) positive neurons in the granule layer of DG is 3,680 (Jinno and Kosaka, 2006). Hippocampome.org reports two neuron types expressing this molecule in that layer: Axo-axonic and SG PV+ Basket cells; moreover, three additional neuron types have undetermined status for this expression: Total Molecular Layer, Outer Molecular Layer, and MOLAX. All other neuron types in the DG granule layer are PV-negative. We interpret these data as indicating that the number of PV positive neurons in the granule layer is *at most* 3,680, and the combined number of PV positive and PV undetermined neurons is *at least* 3,680 (Table 2).

We added weights to each equation based on the experimental method described in the source. We considered unbiased stereology measurements and whole-brain high-throughput image-processing counts as most reliable, because these approaches are specifically designed to obtain accurate population counts. In contrast, we considered electrophysiological ratios and biomarker inequalities relatively less reliable. This is because electrophysiological experiments have comparatively small sample sizes and less emphasis on unbiased sampling, while biomarker inequalities are intrinsically limited by the uncertainty on neuron type with undetermined expression. Therefore, sums from stereology and image-processing were weighted 10:1 against ratios from electrophysiological studies and inequalities from molecular expression data.

Each equation was thus transformed into a normalized, weighted, least squared form and the corresponding residuals were summed into an objective function for each subregion:

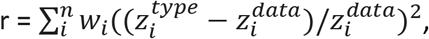

where *r* is the overall residual to be minimized by numerical optimization, *i* is the equation number, *n* is the total number of equations, *w* is the weight, 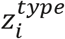 represents the relation between neuron types for the specific equation, and 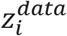 provides the corresponding numerical value extracted from the literature. For instance, in the *ratio* example described above (second column of Table 2), *z^type^* is x_6_/x_8_ (number of CA3 PV+ basket cells divided by number of CA3 axoaxonic cells) and *z^data^* is 1.0333 (31/30). The objective function is then minimized to obtain estimates for the unknown variables (neuron types by layer). Table 1 lists, for each subregion, the number of neuron types, the number of equations generated from the different sources, and the corresponding residual errors. The complete list of equations, interpretations, and sources can be found in the Supplementary Material.

After thorough testing of several optimization algorithms, we adopted PSwarm for minimization of residuals (Vaz and Vicente, 2007). PSwarm is a global optimizer that combines pattern search (Lewis and Torczon, 2002) and particle swarm (Kennedy and Eberhart, 1997) algorithms for bound constrained problems. We used a lower bound of 10 and an upper bound of 10^6^ for each neuron type variable. With the exception of CA2 and CA3, the sets of equations were independent across hippocampal subregions. Thus, DG, CA2+CA3, CA1, Sub, and EC were each optimized separately.

To exploit the stochastic nature of PSwarm, we ran 200 optimization iterations and selected the results with the lowest error. In certain cases, multiple iteration runs tied for lowest error but returned widely differing values for restricted subsets of neuron types. Examination of these situations revealed systematic interdependencies among those neuron types in that their sums were always constant. These relations are indicative of insufficient experimental observations to fully constrain the population sizes of all individual neuron types. For the purpose of subsequent analyses, we assigned an equal population size to each neuron type in the subset. For example, multiple optimization iterations yielded a constant sum of 6,351 for three CA3 neuron types in SO: Interneuron Specific Oriens, O-LMs, and Trilaminars. In this case, the residual error could be minimized by assigning an arbitrary numerical value from 0 to 6351 to any of the three types; then assigning to one of the remaining two types any numerical value from 0 to the difference between 6351 and the value given to the first type; and finally assigning to the third type the difference between 6351 and the sum of values given to the first two types. We assigned a value of 2117 to each of those three neuron types.

We also conducted sensitivity analysis to determine acceptable numerical estimate ranges for every neuron type. Specifically, we computed the lower and upper bounds for each type by programmatically identifying the deviations from the optimal numerical value causing a 5% increase in the residual error for that subregion.

The volumes of the anatomical parcels for density analyses were derived from published data as described in previous quantitative analyses of Hippocampome.org data (Tecuatl et al., 2021). To assess statistical significance of linear correlations, p-values were calculated using Pearson’s coefficient and the number of data points; those p-values were then corrected for multiple testing using the Bonferroni method.

## Results

The primary goal of this work is to obtain estimated population sizes of all neuron types from the six subregions of the rodent hippocampus as defined by Hippocampome.org. To this aim, we apply operations research methods to knowledge extracted from the peer-reviewed scientific literature as recently demonstrated for DG (Attili et al., 2020). Here we improve that approach with the addition of biomarker expression inequalities and expand the effort to the entire hippocampal formation.

The structured literature search described in the Methods yielded a broad collection of 6241 articles, consisting of the union of the 101 ‘core’ articles of Hippocampome.org (hippocampome.org/php/Help_Bibliography.php), 1289 publications cited by the above core articles, and 5035 publications citing the above core articles. We manually triaged this broad collection by perusing titles, abstracts, and illustrations of each publication in search of indications that the content might include data relevant to the quantification of neuron types. This operation resulted in a narrower collection of 570 articles that we mined in-depth by full-text reading. We identified and extracted actionable data from 155 publications, from which we derived 651 equations. Sums from stereological and image processing studies accounted for 186 equations; ratios from electrophysiological studies for 176; and inequalities from molecular biomarker expression for 289 (Table 1). As expected, the highest amount of evidence and, hence, the largest number of equations were available for CA1, which is extensively researched. The least number of articles and subsequently of equations pertained to Sub, with DG, CA3+CA2, and EC falling in between.

In all cases the numerical optimization converged to a relatively small residual error, ranging from 6% in EC to 22% in CA1. These residual errors quantify the extent of inconsistent information in the literature and may be due to a combination of measurement errors, variation in experimental procedures (e.g., animal sex, strain, and age; use of gene or protein expression in biomarker determination; or longitudinal location of the sampled region along the septal-temporal axis of the hippocampus), limit of the scaling rule from mice to rats, and additional factors.

The numerical optimization process yielded the best values and ranges for all 207 variables, each correspond to a Hippocampome.org neuron type in a specific layer, with all numbers referring to rats. In the subsequent analysis, these variables were combined to obtain overall counts by anatomical parcel (subregions and layers), by neuron type (summing across the layers in which they were present), or by broader neuron families with specific properties (e.g., firing pattern or molecular expression). The original values and ranges for all variables, along with the equations and residual errors, are available in the Supplementary Material.

The total number of neurons across all types from all subregions was 2,958,500, with the highest proportion (1,200,734) in DG and the lowest (29,493) in CA2 (Fig. 1A). EC has been divided into the lateral (LEC) and medial (MEC) areas. Surprisingly, LEC has a higher number of neurons at 583,002 compared to the MEC population size at 196,452. Corresponding ranges determined by sensitivity analysis indicated a higher degree of uncertainty for GABAergic than for glutamatergic neurons (Fig. 1B). The total count of interneurons (299,377) was very close to 10% of the overall population, though this proportion varied in individual subregions, from <4% in DG to >25% in MEC. Next, we examined the proportions of excitatory and inhibitory neuron types across the individual layers of each subregion (Fig. 1C). As expected, the majority of glutamatergic cells occupied the principal cell layers. The GABAergic cells displayed a diverse and uneven laminar distribution among subregions, with SO as the most interneuron-rich layer in CA1 and CA3, and SLM in CA2. We also divided the glutamatergic and GABAergic neurons in two families each: principal cells and other glutamatergic cells; and perisomatic and dendritic-targeting interneurons (Fig. 1D). Hippocampome.org does not identify any non-principal glutamatergic cells in CA2 and Sub, nor any dendritic-targeting interneurons in Sub. Among the rest of subregions and families, the proportion of non-principal glutamatergic cells is highest in LEC and lowest in CA1, while the proportion of perisomatic interneurons is highest in MEC and lowest in DG.

**Figure 1:**
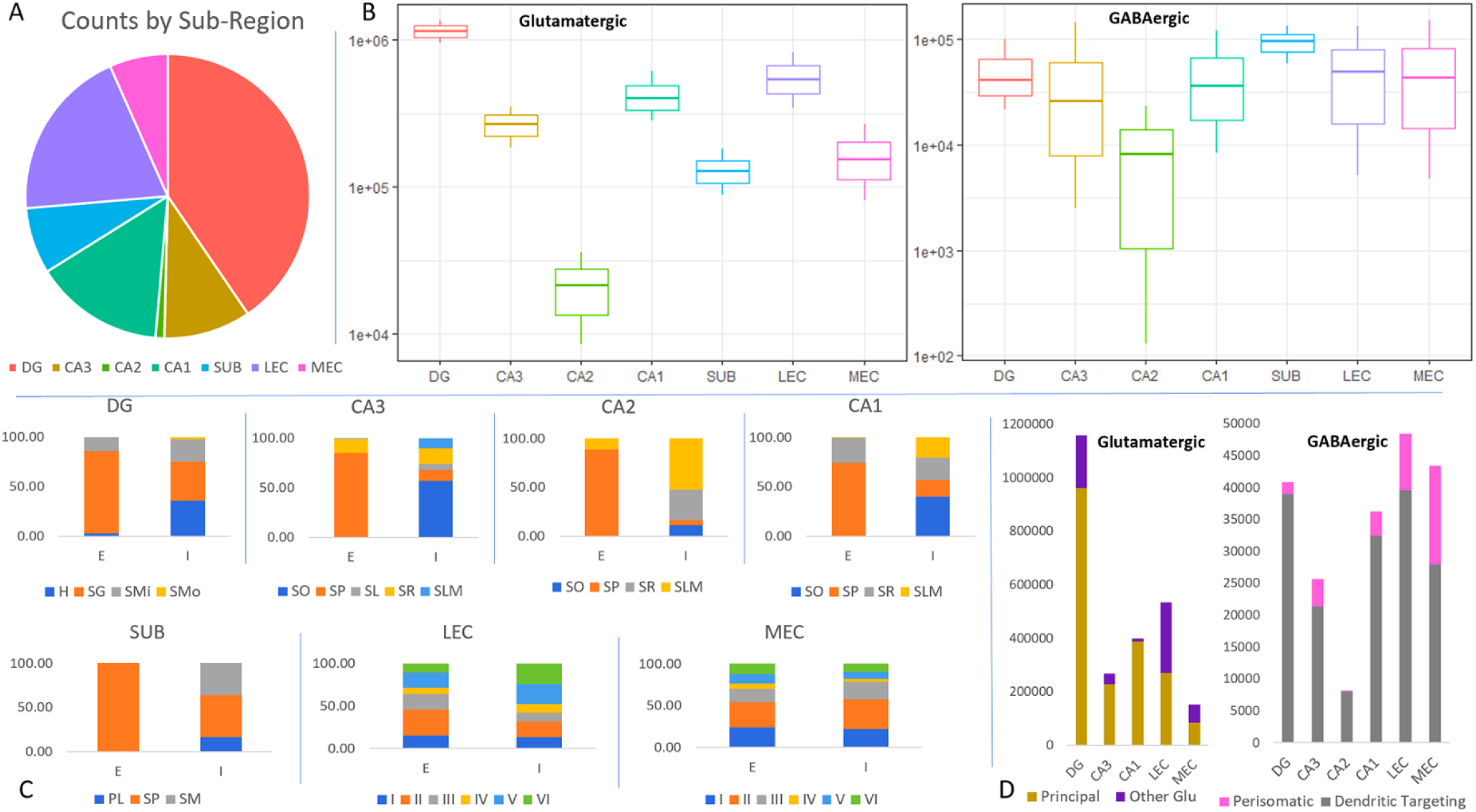
Neuron counts in the rat hippocampal formation by subregion, layer, and neurotransmitter. A. Proportions of neurons by subregions (total counts: DG 1,197,548; CA3 293,278; CA2 29,493; CA1 435,735; SUB 222,992; LEC 583,002; MEC 196,452). B. Totals and ranges (in log scale) for excitatory (left) and inhibitory (right) neurons by sub-region. C. Percentages of neurons across the layers of each subregion for excitatory and inhibitory types. Abbreviations: outer stratum (s.) moleculare (SMo), inner s. moleculare (SMi), s. granulosum (SG), hilus (H); s. lacunosum-moleculare (SLM), s. radiatum (SR), s. lucidum (SL), s. pyramidale (SP), s. oriens (SO); s. moleculare (SM), polymorphic layer (PL); EC layers I–VI. D. Population size by subregion and neuron families: principal cells vs. other glutamatergic cells and dendritic-targeting vs perisomatic cells.

The individual neuron type counts and their upper and lower bound ranges varied widely across subregions (Fig. 2). Generally, dendritic-targeting interneurons were more numerous than perisomatic ones. The most abundant interneuron types in each subregion were DG MOCAPs, CA3 QuadD-LMs, CA2 SLM inhibitory, CA1 Neurogliaforms, SUB SP interneurons, and EC deep layer interneurons. The numerical optimization yielded definite counts and ranges for the majority of neuron types (indicated by blue dots in Fig. 2). These are neuron types that are extensively researched and for which sufficient quantitative evidence is available, including in most cases stereological counts. For approximately a quarter of neuron types (33/122: orange dots in Fig. 2), the lower bound reached 0, making the range significantly wider. These are neuron types for which relevant information is sparser, with often undetermined molecular identity, and typically entangled in numerical interdependencies where the sum of certain subsets is constant. Those numerical interdependencies are also explicitly listed in Table 3. Lastly, four neuron types (pink dot in Fig. 2) returned population size estimates at the lower bound values set by the optimization algorithm (10): DG HIPROMs, DG Outer Molecular Layer interneurons, CA3 Lucidum ORAX, and CA1 Quadrilaminar. These are types whose very existence cannot be reliably confirmed based on available evidence.

**Figure 2:**
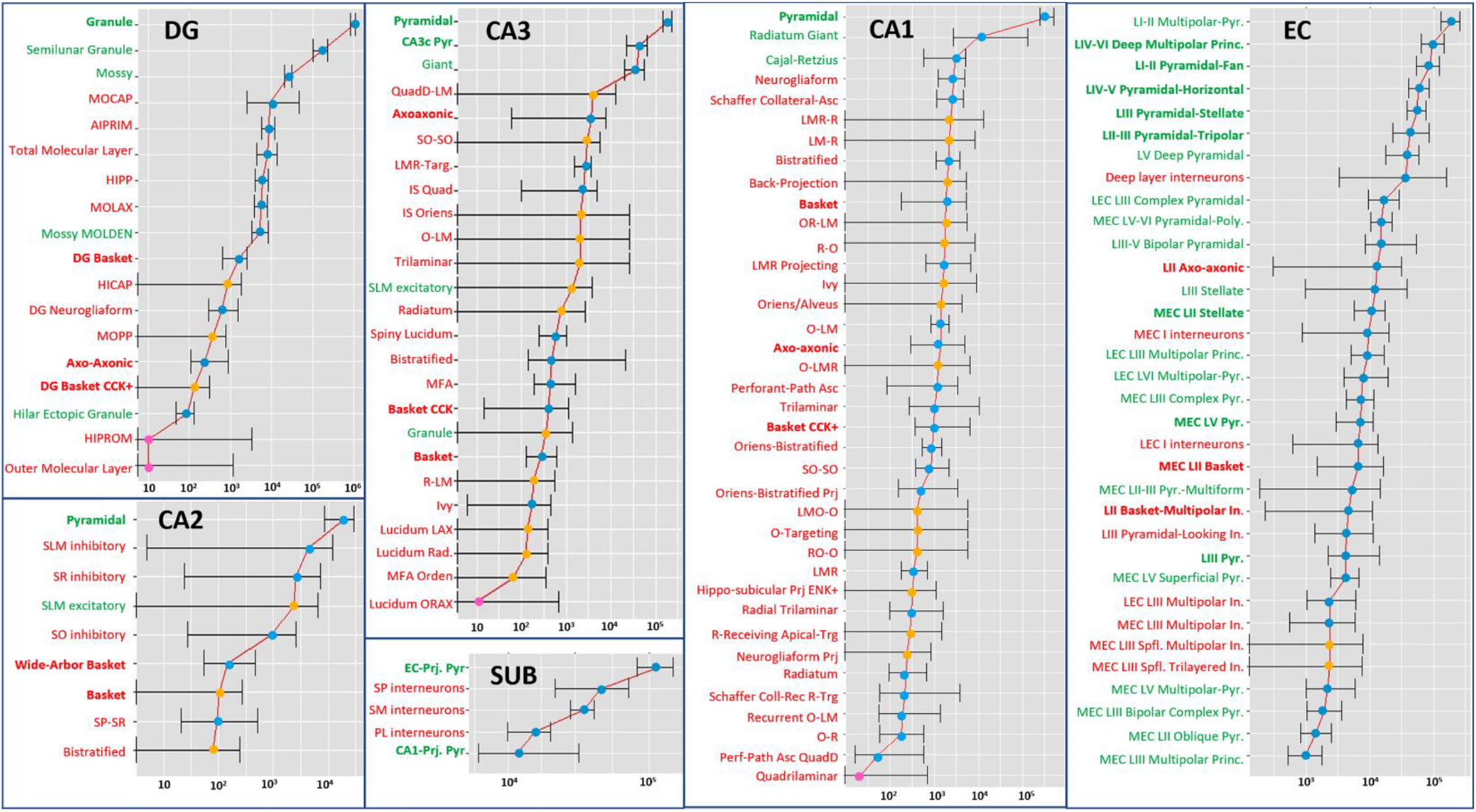
Counts and ranges (in log scale) for all 122 Hippocampome.org neuron types. Excitatory neurons are in green (bold for principal cells), inhibitory in red (bold for perisomatic). Blue dots: higher reliability; orange dots: medium reliability; pink dot: lower reliability.

**Table 3:** Interdependencies among variables and their numerical resolutions.

| Subregion | Variable | Neuron Type | Estimate | Interdependencies |
| --- | --- | --- | --- | --- |
| CA3 | x34 | CA3 IS Oriens SO | 2117 | $x34+x50+x58=6350$<br>Each -> 6350/3 |
|  | x50 | CA3 O-LM SO | 2117 |  |
|  | x58 | CA3 Trilaminar SO | 2117 |  |
| CA1 | x87 | SP IS LMO-O | 90 | $(x88+x95+x98)+x24 = 1962$; $(x88+x95+x98)=981$ & $x24=981$<br>$x91+x92 = 2446$; $x91=2446-x92 = 2446-981 = 1465$<br>$(x87+x94+x97)+x91 = 1734$; $(x87+x94+x97)= 1734-1465 = 269$ |
|  | x88 | SR IS LMO-O | 327 |  |
|  | x91 | SP IS LMR-R | 1465 |  |
|  | x92 | SR IS LMR-R | 981 |  |
|  | x94 | SP IS O-Targeting | 90 |  |
|  | x95 | SR IS O-Targeting | 327 |  |
|  | x97 | SP IS RO-O | 90 |  |
|  | x98 | SR IS RO-O | 327 |  |
| EC | x141 | LI-II Multipolar-Pyramidal LECI | 41397 | $x141+x145=82794$; each -> 82794/2<br>$x152+x164=20781$; each -> 20781/2<br>$x192+x193=4698$; $x192=4698/2=2349$;<br>$x193=4698/2=2349$<br>$x190+x193=8080$; $x190=8080-x193=8080-2349=5731$<br>$x194+x195=5281$; $x194=5281/2=2641$;<br>$x195=5281/2=2641$<br>$x189+x190+x191=19680 \Rightarrow x189+x191=19680-x190=13949$; $x189=13949/2=6975$; $x191=13949/2=6975$<br>$x195+x196+x199=6393 \Rightarrow x196+x199=6393-x195=3752$;<br>$x196=3752/2=1876$; $x199=3752/2=1876$<br>$x196+x198+x199=6738 \Rightarrow x198 = 6738-3752 = 2986$ |
|  | x145 | LI-II Pyramidal-Fan LECI | 41397 |  |
|  | x152 | LII-III Pyramidal-Tripolar LECIII | 10391 |  |
|  | x164 | LIII Stellate LECIII | 10391 |  |
|  | x189 | LII Axo-axonic MECII | 6975 |  |
|  | x190 | LII Axo-axonic LECII | 5731 |  |
|  | x191 | MEC LII Basket | 6975 |  |
|  | x192 | LII Basket-Multipolar Interneuron MECII | 2349 |  |
|  | x193 | LII Basket-Multipolar Interneuron LECII | 2349 |  |
|  | x194 | LEC LIII Multipolar Interneuron | 2641 |  |
|  | x195 | MEC LIII Multipolar Interneuron | 2641 |  |
|  | x196 | MEC LIII Superficial Multipolar Interneuron | 1876 |  |
|  | x198 | LIII Pyramidal-Looking Interneuron | 2986 |  |
|  | x199 | MEC LIII Superficial Trilayered Interneuron | 1876 |  |

Hippocampome.org has collated expression information about ~100 molecular biomarkers for each neuron type (White et al., 2020). Moreover, the knowledge base includes information regarding the intrinsic firing pattern of the neuron types and defines 23 electrophysiological phenotypes accordingly (Komendantov et al., 2019). Given the practical and functional importance of these characterizations, we analyzed the neuron census obtained in this work by grouping the neuron types into molecular and electrophysiological families (Fig. 3). Specifically, we quantified the layer-wise counts for all GABAergic interneurons expressing each of the nineteen most studied biomarkers (Fig. 3A). The highest number of interneurons in the hippocampal formation are positive to Neuropeptide Y, and the lowest number are positive to enkephalin. Among the firing patterns, the majority of neurons in the hippocampal formation was adapting spiking (ASP), while transient stuttering followed by silence (TSTUT.SLN) was exhibited by the least number of neurons (Fig. 3B). Interesting regional and laminar trends also emerged, with delayed non-adapting spiking (D.NASP) almost entirely confined to the DG granular layer and (non-delayed) non-adapting spiking (NASP) more commonly observed in the deep layers of EC.

**Figure 3:**
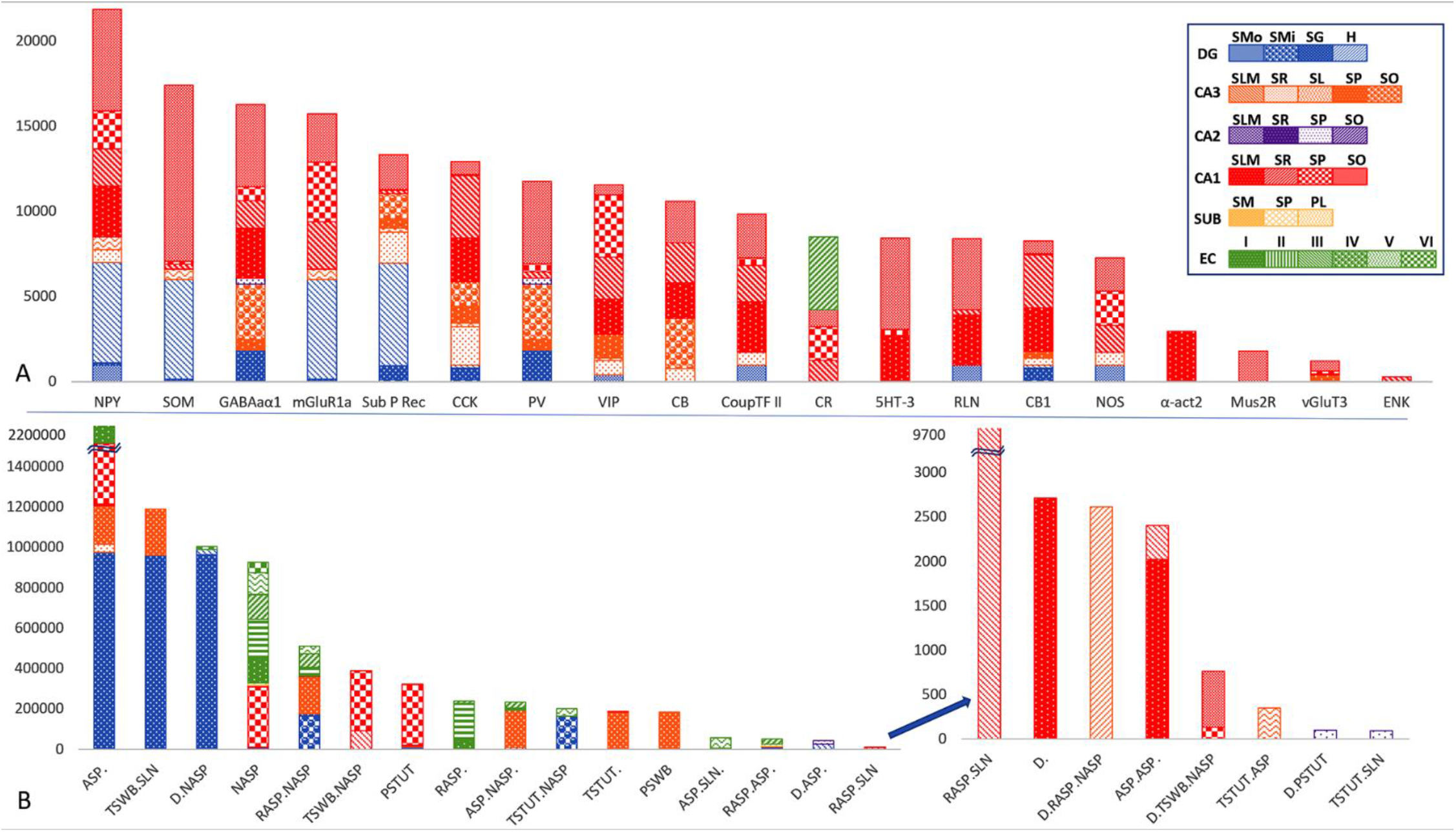
Neuron counts by molecular expression and firing patterns across the 26 layers of the six hippocampal subregions. A. Population sizes of neuron families defined by the expression of 19 molecular biomarkers. CB: Calbindin; CR; Calretinin; PV: Parvalbumin; CB1: Cannabinoid receptor type 1; mGluR1a: metabotropic glutamate receptor 1 alpha; Mus2R: muscarinic type 2 receptor; vGluT3: vesicular glutamate transporter 3; CCK: Cholecystokinin; ENK: Enkephalin; NPY: Neuropeptide Y; SOM: Somatostatin; VIP: Vasoactive intestinal polypeptide; nNOS: Neuronal nitric oxide synthase; RLN: Reelin; 5HT_3 - Serotonin receptor 3; Sub_P_Rec: Substance P receptor; GABA_A_-α_1_: GABA_A_ alpha 1 subunit; a-act2: alpha actinin 2; CoupTF_2: chicken ovalbumin upstream promoter transcription factor IIB. Population sizes of neuron families defined by 23 firing pattern phenotypes. ASP.: adapting spiking; ASP.ASP.: adapting spiking followed by (slower) adapting spiking; ASP.NASP.: non-adapting spiking preceded by adapting spiking; ASP.SLN: silence preceded by adapting spiking; D.: delayed spiking; D.ASP.: delayed adapting spiking; D.RASP.NASP: non-adapting spiking preceded by delayed fast-adapting spiking; D.NASP: delayed non-adapting spiking; D.PSTUT: delayed persistent stuttering; D.TSWB.NASP: non-adapting spiking preceded by delayed transient slow-wave bursting; RASP.: fast-adapting spiking; RASP.ASP.: fast-adapting spiking followed by adapting spiking; RASP.NASP: non-adapting spiking preceded by fast-adapting spiking; RASP.SLM: silence preceded by fast adapting spiking; NASP: non-adapting spiking; PSTUT: persistent stuttering; PSWB: persistent slow-wave bursting; TSTUT.: transient stuttering; TSTUT.ASP.: transient stuttering followed by adapting spiking; TSTUT.NASP: non-adapting spiking preceded by transient stuttering; TSTUT.SLN: silence preceded by transient stuttering; TSWB.NASP: non-adapting spiking preceded by transient slow-wave bursting; TSWB.SLN: silence preceded by transient slow-wave bursting.

Next, we investigated the variation of the population counts for the main excitatory and inhibitory families (principal neurons, other glutamatergic cells, perisomatic interneurons, and dendritic-targeting interneurons) as a function of the volumes of each anatomical parcel (subregion and layer) in the hippocampal formation. If each neuronal population has (approximately) constant density across parcels, the population count should be proportional to the anatomical volumes. If, in contrast, the population counts themselves were approximately constant, then the densities should be inversely proportional to the anatomical volumes. We tested these alternative hypotheses by plotting the population counts against parcel volumes and the densities against the inverse of the volumes for each of the four populations (Fig. 4). As general trends, neuron counts were proportional to anatomical volumes for other (non-principal) glutamatergic neurons and for perisomatic interneurons. In those two cases the densities did not vary with the inverse of volumes. In contrast, in the other populations, principal cells and dendritic-targeting interneurons, the counts did not vary with volumes. Instead, for the dendritic-targeting interneurons, the densities were inversely proportional to the volume with high statical significance (p<10^−4^ after multiple testing correction). The analysis results were the same when analyzing either rat or mouse data. Thus, we conclude that dendritic-targeting interneurons have constant counts independent of the volume of the anatomical parcel in which their somata reside.

**Figure 4:**
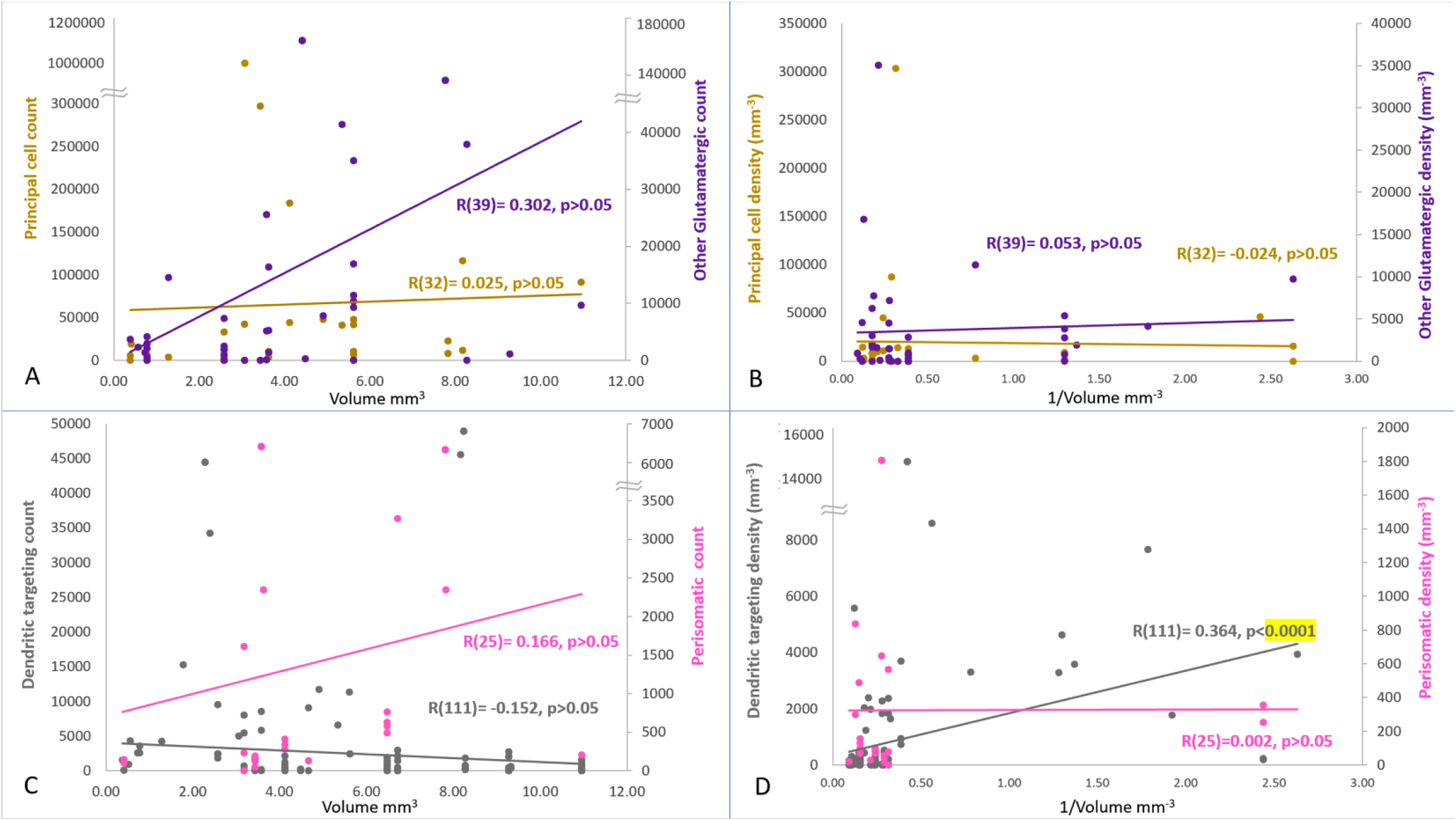
Variation of the counts and densities of four main neuron families by anatomical volumes. Left panels: relations between neuron counts and volumes. Right panels: relations between neuron densities and inverse of volumes. Top panels: principal cells (left axes) and other glutamatergic neurons (right axes). Bottom panels: dendritic-targeting (left axes) and perisomatic (right axes) GABAergic interneurons. The number of neuron types corresponding to each of the 4 families is reported in parentheses after Pearson coefficients.

We have made all software scripts and mined data available through the Supplementary Material for further research and analysis. All neuron type counts (optimization estimates and ranges) have been released on Hippocapome.org on the corresponding neuron pages (Fig. 5). The ‘neuron type census’ page^1^ (hippocampome.org/counts) provides a browsable matrix with the rat and mouse numerical estimates for all 122 neuron types and displays the ranges upon cursor hover-over. As for all Hippocampome.org content, clicking on a value dynamically pulls all underlying experimental evidence supporting the annotated knowledge. In this case, the evidence page includes all literature references and excerpts related to each equation pertaining to that neuron type and corresponding interpretations.

**Figure 5:**
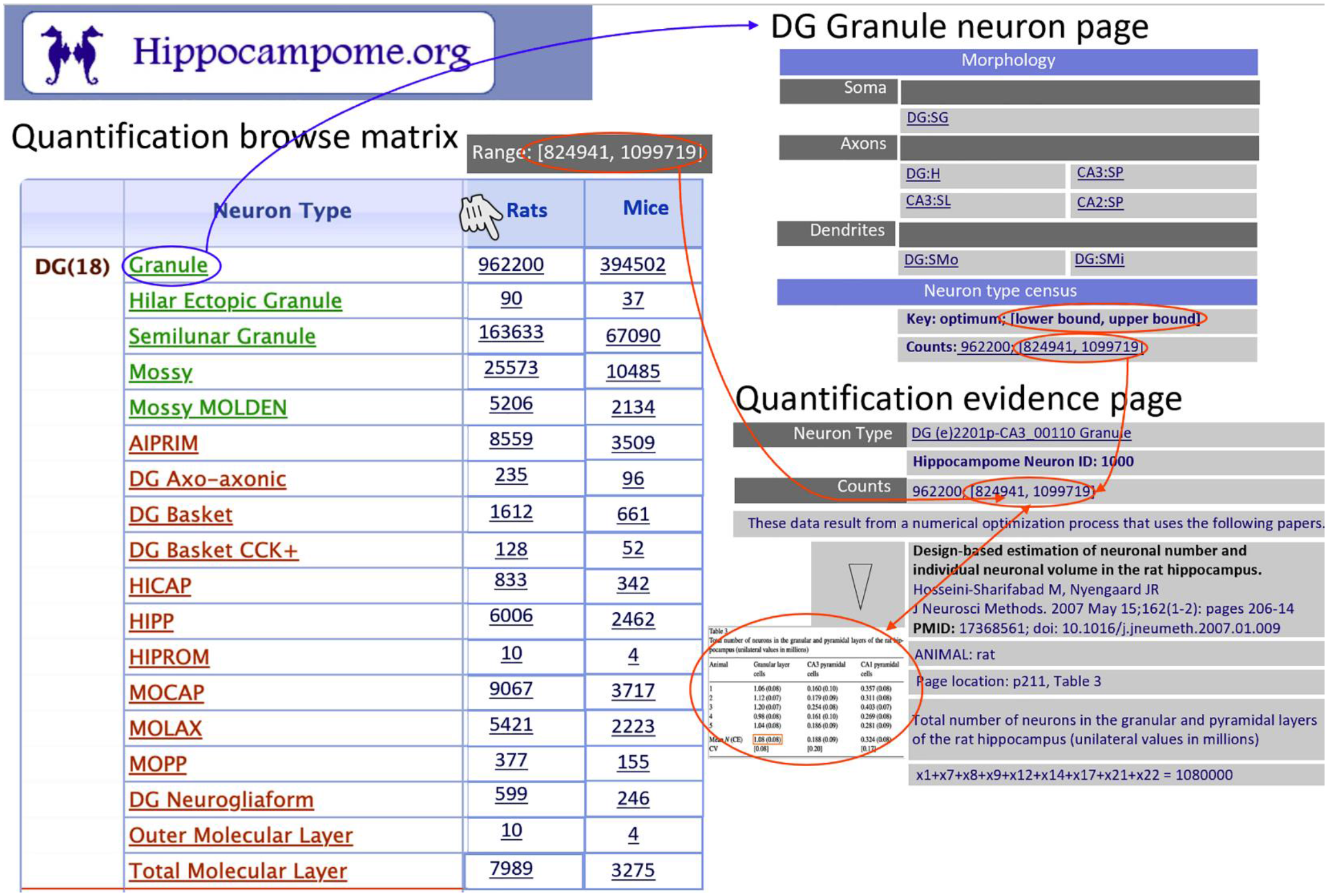
New Hippocampome.org pages and functionality (Hippocampome.org/counts). The neuron type census matrix (left) displays the list of neuron type and their estimated counts for rats and mice with ranges appearing in overlay upon hover-over. The information is also reported for each type in the corresponding neuron page (top right). Every numerical value is dynamically linked to the collection of evidence (bottom right) supporting all equations pertaining to the related neuron type.

The number of neurons in each type can be multiplied with recently derived connection probabilities (Tecuatl et al., 2021) to derive the average numbers of neurons of each type receiving synaptic contacts from a given cell (network divergence) or sending synaptic contacts to a given cell (network convergence). Further multiplying the above values by the average number of computed synaptic contacts per pair of connected cells, also available at Hippocampome.org (Tecuatl et al., 2021) provides an estimate of the total number of synapses a single neuron receives from (or sends to) every neuron type with which it directly interacts in the circuit. For example, the connection probability between CA3 Pyramidal cells is 0.01034, with an average of 7.24 contacts per connected pair. From the count of 183,844 obtained here, it is possible to derive a number of ~13,763 synapses each CA3 Pyramidal cells receives from other *ipsilateral* CA3 Pyramidal cells. This estimate is consistent with the value reported for the quantity of dendritic spines in CA3 Pyramidal cells in strata oriens and radiatum, corresponding to recurrent collaterals (Sunanda et al., 1995, Major et al., 1994). The same quantitative reasoning can be extended to cases for which confirmatory data are not yet available, thus providing a valuable estimate for computational modeling. For instance, it is possible to compute that a CA3 Pyramidal cell receives approximately 792 synapses from CA3 O-LM interneurons in CA3 SLM (connection probability of 0.020 times 18.7 contacts per connected pairs times 2117 presynaptic cells) while an O-LM interneurons receives approximately 4,847 synapses from CA3 Pyramidal cells in stratum oriens (connection probability of 0.0055 times 4.79 contacts per connected pair times 183,844 presynaptic cells).

## Discussion

This work constitutes, to the best of our knowledge, the first comprehensive cell census for any mammalian neural system based on a consistent neuron type classification framework. The systematic application of an efficient numerical optimization workflow to all relevant published evidence yielded population size estimates for all 122 neuron types of Hippocampome.org. Therefore, this open access knowledge base now includes first-approximation cell counts of all known neuron types in the rodent hippocampus and entorhinal cortex defined on the basis of their main neurotransmitters and axonal-dendritic patterns.

Since our method heavily relies on the existing scientific literature, the data mining phase is critically important. We carefully followed a step-by-step literature search process with the goal of obtaining every single published scientific report that was relevant to any of the neuron types of interest. The results clearly reflect the amount of research conducted on specific anatomical regions and cell types. The DG granule and CA1 pyramidal layers, for instance, have been extensively studied by multiple independent teams, hence yielding rich evidence for the corresponding neurons. These mainly consist of glutamatergic principal cells respectively providing the input and output communication pathways of Ammon’s Horn (Wang et al., 2006; Barkai and Hasselmo, 1994; Yang et al., 1996; Hestrin et al., 1990). GABAergic interneurons in those principal cell layers, such as Basket and Axo-axonal cells, providing perisomatic inhibition to the principal cells, are also well characterized compared to the highly diverse interneurons in other layers.

In contrast, the subiculum and CA2, the relatively small area between CA1 and CA3, have been less studied compared to the rest of the hippocampal formation, resulting in fewer constraints for the numerical optimization. Occasionally we found evidence supporting the presence of certain neurons without a corresponding neuron type defined by Hippocampome.org. For example, no excitatory neuron types have been characterized yet in CA3 SLM, but glutamatergic biomarkers were quantified in that layer (Ero et al., 2018). In those cases, we defined a variable for the unknown type(s) and added the relation into the optimization process. By homology with the adjacent area CA1, we speculate that the missing type in this particular instance may be Cajal-Retzius neurons (Quattrocolo and Maccaferri, 2014). Other similar situations likely reflect more complex scenarios, such as the expected diversity of GABAergic interneurons in layers IV, V, and VI of medial and lateral entorhinal cortices for which no detailed identification is available. All ad-hoc variables listed in the supplementary material that do not correspond to identified Hippocampome.org types indicate potential ‘low hanging fruits’ of new neurons awaiting discovery.

Our emphasis on including all relevant published data we could find in the peer-reviewed literature implies that we did not filter any outlier, nor did we attempt to manually reconcile seemingly contrasting data. For example, one stereological study reported 380,000 CA1 principal layer neurons (West et al., 1991), whereas another reported 262,181 (Fitting et al., 2009). Similarly, the ratio between DG SMo MOPP and Neurogliaform cells varies across publications from 6:11 or 0.54 (Armstrong et al., 2011) to 14:17 or 0.82 (Ceranik et al., 1997). The post-optimization residuals for each subregions de facto quantify the overall extent of data variability in the underlying scientific publications. Those values, ranging from 6% to 22% are relatively modest considering that we pooled together mouse and rat data using an admittedly coarse scaling rule (Herculano-Houzel et al., 2006) across a broad heterogeneity of animal strain, sex, caging conditions, and experimental techniques.

In light of this diversity in research methodology, our mathematical optimization approach aimed to find the best estimate accounting for all available data. Moreover, sensitivity analysis allowed us to determine reliability ranges around each estimated optimum value by identifying the upper and lower limits at which the residual for the corresponding subregions would exceed 5% of the minimum. For example, our results indicate an optimal value of 299,408 for the count of Pyramidal neurons in CA1 SP, with a range from 224,807 to 363,841. At those lower and upper bounds, the residual for CA1 increases from the minimum of 22.03% to over 23.13%.

In a limited number of cases, the available evidence was insufficient to uniquely determine an optimal value for the cell counts of a subset of neuron types. These circumstances always resolved to reveal interdependences between neuron types, whereas the number of neurons in each type could vary widely but only as long as the sum over those types would remain constant. These circumstances reflect the inability to unequivocally associate one or more pieces of experimental evidence with a single neuron type. Similar to the situation of ‘unknown’ types described above, these ambiguities identify under-studied areas of research and the need for additional or more specific distinguishing properties between neuron types, especially selective molecular biomarkers.

We analyzed neuron type-specific cell counts by subregion and layer, by transcriptomic expression (White et al., 2020), and by spiking patterns (Komendantov et al., 2019). This mapping of population size estimates to commonly used neuronal properties provides an overview of the proportions of neuron types in each broad anatomical, molecular, or physiological family. Interestingly, the numerical densities of dendritic targeting GABAergic interneurons (but not of perisomatic interneurons, principal cells, or other glutamatergic neurons) were inversely proportional to anatomical volumes. This finding implies that the abundance of these inhibitory cells across the hippocampal formation does not follow the generally observed trend of increasing with the size of the space in which they reside (e.g., Attili et al., 2019). It remains to be established whether this observation extends to other neural systems beyond the hippocampus and entorhinal cortex. Notably, dendritic targeting interneurons are the most diverse group of cells, accounting for half of all Hippocampome.org neuron types.

The complete set of results from this research project are freely released to the scientific community with the publication of this report both as Supplementary Materials and through the searchable and browsable online portal of Hippocampome.org. These include the optimized numerical estimates and ranges for all neuron types, literature excerpts for all supporting evidence and related interpretations, and all associated source code. We anticipate that these data will be used by future projects that involve the development of working models of the rodent brain and related computational simulations. Together with recently reported connection probabilities (Tecuatl et al., 2021), the neuron type population sizes will also enable a more thorough quantification of the hippocampal-entorhinal circuit formation thus extending previous qualitative graph-theoretic analyses (Rees et al., 2016). Last but not least, our methodology can be adapted and expanded to obtain estimated cell counts for the neuron types of other brain regions and animal species.

## Acknowledgements

This project is supported in parts by grants R01NS39600 and U01MH114829. We are grateful to Drs. Padmanabhan Seshaiyer (Professor of Mathematical Sciences, George Mason University) and Siva Venkadesh (from the author’s lab) for critical discussions.

## Author Contributions

SMA performed initial research, methodology generation, data mining, data transformation, numerical optimization, analysis, initial manuscript writing, and generation of tables and figures 1-4. KM contributed to data mining, numerical optimization, and editing of manuscript, figures, and tables. DWW contributed to figure 5 generation and editing of manuscript, figures, and tables. Both KM and DWW provided their subject matter expertise during critical discussions. GAA contributed to methodology generation, data interpretation, analysis design, manuscript writing, and editing of figures and tables, obtained and administered research funding, and supervised all aspects of the project.

## Data Availability Statement

The data that supports the findings of this study are available in the supplementary material of this article.

Supplementary Material file contains variables, equations, evidence, numerical ranges, individual results from multiple optimization iterations, and source code. The accompanying ‘Read Me’ file describes the organization of the provided data.

## Footnotes

Access to all data and functionality described in the manuscript are provided via hippocampome.org/counts with a password. Please for the access code; the password protection and this note will be removed upon acceptance of the manuscript in a peer-reviewed journal.

