## SupplementaryData for "Quantification of neuron types in the rodent hippocampal formation by data mining and numerical optimization": ReadMe.txt

The supplementary data file is an excel document containing 25 tabs. There are 5 tabs for each subregion for - Equations, Biomarkers, Results, Manual and Iterations.

Equations - Data from the 5 five types of sources - stereological, computational, electrophysiological, biomakers, modeling and others has been extracted. These data are used to compute linear equations which are transformed in normalized format for optimization. This tab contains these equations, their source names, interpretation and other related information.

Biomarkers - This tabs contains the steps involved in the data transformation of the biomarker results from the literature. Data from scientific literature is converted into rat neuronal numbers for usage.

Results - This tab contains the variables assigned to each neuron type/group, optimum values generated by the algorithm and the lower/upper bounds calculated. Multiple iterations were run and the ones with lowest error were used to report the results. The ranges are the result of our sensitivity analysis. Please refer to methods and results section for more details on the computation of these values.

Manual - The residual errors for each equation was examined. Column A lists the neuron name, Column B - variable name, Column C - Optimum neuron counts generated by the algorithm, Column D - residual errors, Column E - weights, Column F - neuron counts entered by user for testing, Column G - residual errors, Column H - weights, Column I - corresponding equations.

Iterations - For each subregion, we ran 200 iterations of our numerical optimization algorithm and chose the iterations with the lowest error value. The data presented in this tab are the iterations that generated the lowest error value. Trends between neuron type quantities were studied using these data.

The 'Source-Code' folder contains 5 files for the 6 subregions. CA2 and CA3 are combined into one file. These files contain the code for running the numerical optimization algorithm. The program is written to generate 200 iterations and write the results to a file.
