## SupplementaryData for "Quantification of neuron types in the rodent hippocampal formation by data mining and numerical optimization": CA1.R

```
# Define a list with the problem definition
rm(Problem, outputfcn, Options, Result)
Problem <- list()
Problem$Variables <- 64
Problem$objf <- function(x) {
  ##################################
    #Lee
    ((((58*x[82])/(46*x[124]) - 1)^2)*1) +
    #Kawaguchi
    ((((11*(x[75]+x[79]))/(9*(x[121]+x[122]+x[124]+x[125])) - 1 )^2)*1) +
    #Mcbain
    ((((36*x[111])/(15*(x[108]+x[109]+x[110])) - 1 )^2)*1) +
    ((((4*x[111])/(15*x[128]) - 1 )^2)*1) +
    #Sik
    ((((x[78]+x[79]+x[80]+x[81]+x[82])/(6*(x[74]+x[75])) - 1 )^2)*1) +
    ((((2*(x[74]+x[75]))/(x[85]+x[86]) - 1 )^2)*1) +
    (((x[77]/(x[85]+x[86]) - 1 )^2)*1) +
    #Buhl
    ((((11*(x[79]+x[81]))/(31*x[86]) - 1 )^2)*1) +
    (((x[86]/x[75] - 1 )^2)*1) +
    #Hajos
    (((x[108]/(7*x[86]) - 1 )^2)*1) +
    ((((2*x[86])/x[132] - 1 )^2)*1) +
    ((((3*x[132])/(2*x[102]) - 1 )^2)*1) +
    (((x[102]/x[121]  - 1 )^2)*1) +
    (((x[121]/x[93] - 1 )^2)*1) +
    (((x[93]/(3*x[118]) - 1 )^2)*1) +
    (((x[118]/(x[104]+x[105]) - 1 )^2)*1) +
    #Ali
    (((x[108]/(7*x[109]) - 1 )^2)*1) +
    #Vida
    ((((5*x[82])/(2*(x[124]+x[125])) - 1 )^2)*1) +
    ((((3*(x[124]+x[125]))/(5*(x[116]+x[117])) - 1 )^2)*1) +
    ((((x[116]+x[117])/(3*(x[102]+x[103])) - 1 )^2)*1) +
    #Svoboda
    ((((17*(x[74]+x[78]+x[80]))/(12*(x[108]+x[109])) - 1 )^2)*1) +
    ((((22*(x[108]+x[109]))/(17*x[112]) - 1 )^2)*1) +
    ((((24*x[112])/(22*x[130]) - 1 )^2)*1) +
    #Katona
    (((x[108]/(3*x[113]) - 1 )^2)*1) +
    #Cope
    ((((7*x[82])/(8*x[124]) - 1 )^2)*1) +
    #Maccaferri
    ((((7*x[108])/(8*x[112]) - 1 )^2)*1) +
    ((((9*x[112])/(7*x[83]) - 1 )^2)*1) +
    ((((2*x[83])/(3*x[74]) - 1 )^2)*1) +
    #Cossart2000
    (((3*x[108]/(5*(x[116]+x[117])) - 1 )^2)*1) +
    ((((3*(x[74]+x[75]+x[76]+x[78]+x[79]+x[80]+x[81]+x[82]+x[83]+x[84]))/(4*x[85]) - 1 )^2)*1) +
    ((((x[116]+x[117])/x[85] - 1 )^2)*1) +
    ((((5*x[115])/x[108] - 1 )^2)*1) +
    (((x[132]/x[130] - 1 )^2)*1) +
    #Klausberger
    ((((4*x[83])/(x[104]+x[105]) - 1 )^2)*1) +
    #Mercer
    ((((x[79]+x[81])/(2*x[86]) - 1 )^2)*1) +
    #Cossart98
    ((((2*(x[78]+x[79]+x[80]+x[81]+x[82]))/(7*x[74]) - 1 )^2)*1) +
    ((((3*x[74])/(x[85]+x[86]) - 1 )^2)*1) +
    ((((x[85]+x[86])/(2*x[128]) - 1 )^2)*1) +
    ((((7*x[128])/(3*x[108]) - 1 )^2)*1) +
    ((((15*x[108])/(7*(x[124]+x[125])) - 1 )^2)*1) +
    ((((x[124]+x[125])/(3*(x[116]+x[117])) - 1 )^2)*1) +
    #Kogo
    ((((3*x[108])/(13*x[112]) - 1 )^2)*1) +
    #Glickfield
    ((((26*(x[80]+x[81]+x[82]))/(29*(x[78]+x[79])) - 1 )^2)*1) +
    #F
    (((x[107]/x[82] - 1 )^2)*1) +
    (((x[82]/(2*(x[104]+x[105])) - 1 )^2)*1) +
    #Daw
    ((((27*(x[81]+x[82]))/(26*x[86]) - 1 )^2)*1) +
    ((((19*x[86])/(27*(x[131]+x[132])) - 1 )^2)*1) +
    ((((12*(x[131]+x[132]))/(19*x[79]) - 1 )^2)*1) +
    #T
    ((((35*(x[99]+x[100]+x[101]))/(13*(x[106]+x[107])) - 1 )^2)*1) +
    #Ali98_2
    ((((8*(x[78]+x[79]+x[80]+x[81]+x[82]))/(9*(x[85]+x[86]+x[130])) - 1 )^2)*1) +
    #E
    ((((69*x[82])/(72*x[127]) - 1 )^2)*1) +
    #Elfant
    ((((5*x[103])/(2*x[122]) - 1 )^2)*1) +
    ((((9*x[122])/(10*x[117]) - 1 )^2)*1) +
    #Z
    ((((26*x[112])/(24*x[108]) - 1 )^2)*1) +
    ((((11*(x[75]+x[79]+x[86]))/(20*(x[108]+x[110])) - 1 )^2)*1) +
    ((((4*(x[108]+x[110]))/(11*x[112]) - 1 )^2)*1) +
    ((((3*x[112])/(4*(x[102]+x[103])) - 1 )^2)*1) +
    ((((2*(x[102]+x[103]))/(3*x[106]) - 1 )^2)*1) +
    ((((2*x[106])/(x[124]+x[125]) - 1 )^2)*1) +
    ((((5*(x[124]+x[125]))/(4*(x[116]+x[117]+x[130])) - 1 )^2)*1) +
    #Varga
    ((((7*(x[78]+x[79]+x[83]))/(12*(x[74]+x[75])) - 1 )^2)*1) +
    ((((8*(x[74]+x[75]))/(7*(x[85]+x[86]+x[112])) - 1 )^2)*1) +
    #Pawelzik
    ((((4*x[79])/(25*x[78]) - 1 )^2)*1) +
    (((x[86]/(4*x[85]) - 1 )^2)*1) +
    ((((6*x[79])/(25*x[81]) - 1 )^2)*1) +
    (((x[81]/(2*x[80]) - 1 )^2)*1) +
    ((((2*x[81])/(3*x[82]) - 1 )^2)*1) +
    (((x[132]/x[128] - 1 )^2)*1) +
    ((((5*(x[119]+x[120]))/(2*(x[124]+x[125])) - 1 )^2)*1) +
    ((((5*(x[119]+x[120]))/(2*(x[116]+x[117])) - 1 )^2)*1) +
    (((x[78]/(4*x[128]) - 1 )^2)*1) +
    #tyan
    ((((5*x[108])/(18*x[113]) - 1 )^2)*1) +
    ((((3*x[113])/(5*x[83]) - 1 )^2)*1) +
    ((((2*x[83])/x[128] - 1 )^2)*1) +
    #Ganter
    ((((57*x[76])/(4*(x[76]+x[77]+x[83]+x[108]+x[109]+x[111]+x[112]+x[126]+x[128]+x[130]+x[93]+x[110]+x[113])) - 1 )^2)*1) +
    #ABA
    ((((x[71]+x[90]+x[103]+x[105]+x[106]+x[107]+x[117]+x[120]+x[122]+x[125])/87924 - 1 )^2)*10) +
    ((((x[70]+x[73]+x[82]+x[88]+x[89]+x[92]+x[95]+x[98]+x[101]+x[102]+x[104]+x[115]+x[116]+x[118]+x[119]+x[121]+x[123]+x[124]+x[127]+x[129]+x[132])/88798 - 1 )^2)*10) +
    ((((x[74]+x[76]+x[77]+x[78]+x[80]+x[83]+x[85]+x[93]+x[99]+x[108]+x[109]+x[110]+x[111]+x[112]+x[113]+x[126]+x[128]+x[130])/77227 - 1 )^2)*10) +
    #Ero
    (((x[71]/54284 - 1)^2)*1) +
    ((((x[90]+x[103]+x[105]+x[106]+x[107]+x[117]+x[120]+x[122]+x[125])/57983 - 1 )^2)*1) +
    ((((x[74]+x[76]+x[77]+x[78]+x[80]+x[83]+x[85]+x[93]+x[99]+x[108]+x[109]+x[110]+x[111]+x[112]+x[113]+x[126]+x[128]+x[130])/6710 - 1 )^2)*1) +
    #new var
    #(((p[65]/106593 - 1)^2)*1) +
    #w/so e added
    ((((x[69]+x[72])/630767 - 1 )^2)*1) +
    ((((x[70]+x[73])/166551 - 1)^2)*1) +
    ((((x[75]+x[79]+x[81]+x[84]+x[86]+x[87]+x[91]+x[94]+x[96]+x[97]+x[100]+x[114]+x[131])/5310 - 1 )^2)*1) +
    ((((x[82]+x[88]+x[89]+x[92]+x[95]+x[98]+x[101]+x[102]+x[104]+x[115]+x[116]+x[118]+x[119]+x[121]+x[123]+x[124]+x[127]+x[129]+x[132])/27750 - 1 )^2)*1) +
    #M
    ((((x[71]+x[90]+x[103]+x[105]+x[106]+x[107]+x[117]+x[120]+x[122]+x[125])/163988 - 1 )^2)*10) +
    ((((x[74]+x[76]+x[77]+x[78]+x[80]+x[83]+x[85]+x[93]+x[99]+x[108]+x[109]+x[110]+x[111]+x[112]+x[113]+x[126]+x[128]+x[130])/114861 - 1 )^2)*10) +
    ((((x[69]+x[72]+x[75]+x[79]+x[81]+x[84]+x[86]+x[87]+x[91]+x[94]+x[96]+x[97]+x[100]+x[114]+x[131])/376950 - 1)^2)*10) +
    ((((x[70]+x[73]+x[82]+x[88]+x[89]+x[92]+x[95]+x[98]+x[101]+x[102]+x[104]+x[115]+x[116]+x[118]+x[119]+x[121]+x[123]+x[124]+x[127]+x[129]+x[132])/148021 - 1 )^2)*10) +
    #Bezaire numbers
    (((x[69]/311500 - 1 )^2)*10) +
    ((((x[85]+x[86])/2210 - 1 )^2)*10) +
    (((x[106]/3580 - 1 )^2)*10) +
    ((((x[124])/400 - 1)^2)*10) +
    ((((x[80]+x[81]+x[82])/3600 - 1 )^2)*10) +
    ((((x[99]+x[100]+x[101])/8810 - 1 )^2)*1) +
    (((x[108]/1640 - 1 )^2)*10) +
    ((((x[74]+x[75])/1470 - 1 )^2)*10) +
    ((((x[78]+x[79])/5530 - 1 )^2)*1) +
    (((x[117]/490 - 1)^2)*10) +
    ((((x[87]+x[88]+x[94]+x[95]+x[97]+x[98])/1250 - 1)^2)*1) +
    ((((x[89]+x[90])/1970 - 1)^2)*1) +
    #
    ((((x[69]+x[72]+x[75]+x[79]+x[81]+x[84]+x[86]+x[87]+x[91]+x[94]+x[96]+x[97]+x[100]+x[114]+x[131])/387000 - 1)^2)*10) +
    ((((x[69]+x[72]+x[75]+x[79]+x[81]+x[84]+x[86]+x[87]+x[91]+x[94]+x[96]+x[97]+x[100]+x[114]+x[132])/380000 - 1 )^2)*10) +
    ((((x[69]+x[72]+x[75]+x[79]+x[81]+x[84]+x[86]+x[87]+x[91]+x[94]+x[96]+x[97]+x[100]+x[114]+x[131])/324000 -1 )^2)*10) +
    ((((x[69]+x[72]+x[75]+x[79]+x[81]+x[84]+x[86]+x[87]+x[91]+x[94]+x[96]+x[97]+x[100]+x[114]+x[131])/382000 - 1 )^2)*10) +
    ((((x[69]+x[72]+x[75]+x[79]+x[81]+x[84]+x[86]+x[87]+x[91]+x[94]+x[96]+x[97]+x[100]+x[114]+x[131])/650000 - 1 )^2)*10) +
    ((((x[69]+x[72]+x[75]+x[79]+x[81]+x[84]+x[86]+x[87]+x[91]+x[94]+x[96]+x[97]+x[100]+x[114]+x[131])/323300 - 1 )^2)*10) +
    ((((x[69]+x[72]+x[75]+x[79]+x[81]+x[84]+x[86]+x[87]+x[91]+x[94]+x[96]+x[97]+x[100]+x[114]+x[131])/375000 - 1 )^2)*10) +
    ((((x[69]+x[72]+x[75]+x[79]+x[81]+x[84]+x[86]+x[87]+x[91]+x[94]+x[96]+x[97]+x[100]+x[114]+x[131])/346842 - 1 )^2)*10) +
    ((((x[69]+x[72]+x[75]+x[79]+x[81]+x[84]+x[86]+x[87]+x[91]+x[94]+x[96]+x[97]+x[100]+x[114]+x[131])/262181 - 1 )^2)*10) +
    ((((x[69]+x[72]+x[75]+x[79]+x[81]+x[84]+x[86]+x[87]+x[91]+x[94]+x[96]+x[97]+x[100]+x[114]+x[131])/425000 - 1 )^2)*10) +
    ((((x[69]+x[72]+x[75]+x[79]+x[81]+x[84]+x[86]+x[87]+x[91]+x[94]+x[96]+x[97]+x[100]+x[114]+x[131])/200000 - 1 )^2)*10) +
    ((((x[69]+x[72]+x[75]+x[79]+x[81]+x[84]+x[86]+x[87]+x[91]+x[94]+x[96]+x[97]+x[100]+x[114]+x[131])/195122 - 1 )^2)*10) +
    #JinnoKosaka
    ((((x[69]+x[72])/575549 - 1)^2)*10) +
    #Coulin
    ((((x[69]+x[70]+x[71]+x[72]+x[73]+x[74]+x[75]+x[76]+x[77]+x[78]+x[79]+x[80]+x[81]+x[82]+x[83]+x[84]+x[85]+x[86]+x[87]+x[88]+x[89]+x[90]+x[91]+x[92]+x[93]+x[94]+x[95]+x[96]+x[97]+x[98]+x[99]+x[100]+x[101]+x[102]+x[103]+x[104]+x[105]+x[106]+x[107]+x[108]+x[109]+x[110]+x[111]+x[112]+x[113]+x[114]+x[115]+x[116]+x[117]+x[118]+x[119]+x[120]+x[121]+x[122]+x[123]+x[124]+x[125]+x[126]+x[127]+x[128]+x[129]+x[130]+x[131]+x[132])/2056098 - 1)^2)*10) +
    ((((x[71]/2756) - 1)^2)*10) +
    #Poolos
    ((((x[69]+x[70])/(40*(x[72]+x[73])) - 1)^2)*1) +
    #F.P
    ((max(0,(x[129]/(0.11*(x[113]+x[116]+x[117]+x[130])) - 1))^2)*1) +
    #Kosaka PV+
    ((max(0,((x[74]+ x[76]+ x[78]+ x[85]+ x[108])/(0.26*(x[74]+x[76]+x[77]+x[78]+x[80]+x[83]+x[85]+x[93]+x[99]+x[108]+x[109]+x[110]+x[111]+x[112]+x[113]+x[126]+x[128]+x[130])) - 1))^2)*1) +
    ((max(0,(1 - (x[74]+ x[76]+ x[78]+ x[85]+ x[108]+x[83]+x[112])/(0.26*(x[74]+x[76]+x[77]+x[78]+x[80]+x[83]+x[85]+x[93]+x[99]+x[108]+x[109]+x[110]+x[111]+x[112]+x[113]+x[126]+x[128]+x[130]))))^2)*1) +
    ((max(0,((x[75]+ x[79]+ x[86])/(0.48*(x[75]+x[79]+x[81]+x[84]+x[86]+x[87]+x[91]+x[94]+x[96]+x[97]+x[100]+x[114]+x[131])) - 1))^2)*1) +
    ((max(0,(1 - (x[75]+ x[79]+ x[86]+x[84]+x[131]+x[114])/(0.48*(x[75]+x[79]+x[81]+x[84]+x[86]+x[87]+x[91]+x[94]+x[96]+x[97]+x[100]+x[114]+x[131]))))^2)*1) +
    ((max(0,((x[118]+ x[119]+ x[123])/(0.03*(x[82]+x[88]+x[89]+x[92]+x[95]+x[98]+x[101]+x[102]+x[104]+x[115]+x[116]+x[118]+x[119]+x[121]+x[123]+x[124]+x[127]+x[129]+x[132])) - 1))^2)*1) +
    ((max(0,(1 - (x[118]+ x[119]+ x[123]+x[115]+x[132])/(0.03*(x[82]+x[88]+x[89]+x[92]+x[95]+x[98]+x[101]+x[102]+x[104]+x[115]+x[116]+x[118]+x[119]+x[121]+x[123]+x[124]+x[127]+x[129]+x[132]))))^2)*1) +
    #Li
    ((max(0,((x[74]+x[75])/2000 - 1))^2)*1) +
    ((max(0,(1 - (x[74]+x[75])/500))^2)*1) +
    #Jinno
    ((max(0,((x[74]+ x[76]+ x[78]+ x[85]+ x[108])/3677 - 1 ))^2)*1) +
    ((max(0,(1 - (x[74]+ x[76]+ x[78]+ x[85]+ x[108]+x[83]+x[112])/3677 ))^2)*1) +
    ((max(0,((x[75]+ x[79]+ x[86])/11192 - 1 ))^2)*1) +
    ((max(0,(1 - (x[75]+ x[79]+ x[86]+x[84]+x[131]+x[114])/11192 ))^2)*1) +
    ((max(0,((x[118]+ x[119]+ x[123])/466 - 1 ))^2)*1) +
    ((max(0,(1 - ( x[118]+ x[119]+ x[123]+x[132]+x[115])/466 ))^2)*1) +
    ((max(0,((x[120])/85 - 1 ))^2)*1) +
    ((max(0,((x[77]+ x[85]+ x[108]+ x[109]+ x[110]+ x[111]+ x[112]+ x[113]+ x[130])/8122 - 1 ))^2)*1) +
    ((max(0,(1 - (x[77]+ x[85]+ x[108]+ x[109]+ x[110]+ x[111]+ x[112]+ x[113]+ x[130]+x[99])/8122 ))^2)*1) +
    ((max(0,((x[86]+x[114])/1323 - 1 ))^2)*1) +
    ((max(0,(1 - (x[86]+x[87]+ x[91]+ x[94]+ x[97]+ x[100])/1323 ))^2)*1) +
    ((max(0,((x[102]+ x[118]+ x[119]+ x[121]+x[115])/311 - 1 ))^2)*1) +
    ((max(0,(1 - (x[102]+ x[118]+ x[119]+ x[121]+x[88]+ x[92]+ x[95]+ x[98]+ x[101]+ x[129])/311 ))^2)*1) +
    ((max(0,((x[103]+x[120]+x[122])/43 - 1 ))^2)*1) +
    ((max(0,(1 - (x[103]+x[120]+x[122]+x[106]+x[107])/43 ))^2)*1) +
    ((max(0,((x[83]+ x[126])/631 - 1 ))^2)*1) +
    ((max(0,(1 - ( x[83]+ x[126]+x[80])/631 ))^2)*1) +
    ((max(0,((x[84]+ x[87]+ x[91]+ x[94]+ x[96]+ x[97])/6053 - 1 ))^2)*1) +
    ((max(0,(1 - (x[84]+ x[87]+ x[91]+ x[94]+ x[96]+ x[97]+x[81]+x[114])/6053 ))^2)*1) +
    ((max(0,((x[88]+ x[89]+ x[92]+ x[95]+ x[98]+ x[102]+ x[127])/1639 - 1 ))^2)*1) +
    ((max(0,(1 - (x[88]+ x[89]+ x[92]+ x[95]+ x[98]+ x[102]+ x[127]+x[82]+x[115])/1639 ))^2)*1) +
    ((max(0,((x[90]+x[103])/2500 - 1 ))^2)*1) +
    ((max(0,((x[93]+ x[112]+ x[126])/841 - 1 ))^2)*1) +
    ((max(0,((x[87]+ x[94]+ x[96]+ x[97])/5385 - 1 ))^2)*1) +
    ((max(0,(1 - (x[87]+ x[94]+ x[96]+ x[97]+x[91]+x[114])/5385 ))^2)*1) +
    ((max(0,((x[88]+ x[95]+ x[98]+ x[118]+ x[119]+ x[127])/1708 - 1 ))^2)*1) +
    ((max(0,(1 - (x[88]+ x[95]+ x[98]+ x[118]+ x[119]+ x[127]+x[92]+x[115])/1708 ))^2)*1) +
    ((max(0,((x[120])/2572 - 1 ))^2)*1) +
    ((max(0,(1 - (x[120]+x[106]+x[107])/2572 ))^2)*1) +
    ((max(0,((x[109]+ x[111]+ x[126]+ x[128])/6659 - 1 ))^2)*1) +
    ((max(0,(1 - (x[109]+ x[111]+ x[126]+ x[128]+ x[99]+x[108]+x[130])/6659 ))^2)*1) +
    ((max(0,(1 - (x[100]+x[114])/2482 ))^2)*1) +
    ((max(0,((x[104]+ x[121]+ x[124]+ x[127])/2140 - 1 ))^2)*1) +
    ((max(0,(1 - (x[104]+ x[121]+ x[124]+ x[127]+x[101]+x[115])/2140 ))^2)*1) +
    ((max(0,((x[105]+ x[122]+ x[125])/3950 - 1 ))^2)*1) +
    ((max(0,((x[77]+ x[85]+ x[99]+ x[108]+ x[109]+ x[128])/6924 - 1 ))^2)*1) +
    ((max(0,(1 - (x[77]+ x[85]+ x[99]+ x[108]+ x[109]+ x[128]+x[126])/6924 ))^2)*1) +
    ((max(0,((x[86]+ x[100]+ x[131])/8780 - 1 ))^2)*1) +
    ((max(0,(1 - (x[86]+ x[100]+ x[131]+x[114])/8780 ))^2)*1) +
    ((max(0,((x[101]+ x[102]+ x[118]+ x[132])/2002 - 1 ))^2)*1) +
    ((max(0,(1 - (x[101]+ x[102]+ x[118]+ x[132]+x[121]+ x[123]+ x[127]+x[115])/2002 ))^2)*1) +
    ((max(0,((x[103]+ x[106]+ x[107])/3566 - 1 ))^2)*1) +
    ((max(0,(1 - (x[103]+ x[106]+ x[107]+x[122])/3566 ))^2)*1) +
    ((max(0,((x[80])/777 - 1 ))^2)*1) +
    ((max(0,(1 - (x[80]+x[83]+x[85])/777 ))^2)*1) +
    ((max(0,((x[81]+x[131])/2154 - 1 ))^2)*1) +
    ((max(0,(1 - (x[81]+x[131]+x[84]+x[86]+x[114])/2154 ))^2)*1) +
    ((max(0,((x[82]+ x[102]+ x[104]+ x[116]+ x[119]+ x[121]+ x[123]+ x[124]+ x[132])/1915 - 1 ))^2)*1) +
    ((max(0,(1 - (x[82]+ x[102]+ x[104]+ x[116]+ x[119]+ x[121]+ x[123]+ x[124]+p[64 ]+x[115])/1915 ))^2)*1) +
    ((max(0,((x[103]+ x[105]+ x[117]+ x[120]+ x[122]+ x[125])/1179 - 1 ))^2)*1) +
    ((max(0,(1 - (x[103]+ x[105]+ x[117]+ x[120]+ x[122]+ x[125]+x[106]+x[107])/1179 ))^2)*1) +
    ((max(0,((x[77]+x[99])/1857 - 1 ))^2)*1) +
    ((max(0,(1 - (x[77]+x[99]+x[126])/1857))^2)*1) +
    ((max(0,((x[96]+ x[100])/7809 - 1 ))^2)*1) +
    ((max(0,(1 - (x[96]+ x[100]+x[114])/7809 ))^2)*1) +
    ((max(0,((x[101])/2519 - 1 ))^2)*1) +
    ((max(0,(1 - (x[101]+x[102]+ x[121]+ x[123]+ x[127]+x[115])/2519 ))^2)*1) +
    ((max(0,(1 - (x[103]+ x[106]+ x[107]+ x[122])/5399 ))^2)*1) +
    #Kim
    ((max(0,((x[74]+ x[76]+ x[78]+ x[85]+ x[108])/3915 - 1 ))^2)*1) +
    ((max(0,(1 - (x[74]+ x[76]+ x[78]+ x[85]+ x[108]+x[83]+x[112])/3915 ))^2)*1) +
    ((max(0,((x[75]+ x[79]+ x[86])/4732 - 1 ))^2)*1) +
    ((max(0,(1 - (x[75]+ x[79]+ x[86]+x[84]+x[131])/4732 ))^2)*1) +
    ((max(0,((x[118]+ x[119]+ x[123])/870 - 1 ))^2)*1) +
    ((max(0,(1 - (x[118]+ x[119]+ x[123]+x[132])/870 ))^2)*1) +
    ((max(0,((x[120])/44 - 1 ))^2)*1) +
    ((max(0,((x[77]+ x[85]+ x[108]+ x[109]+ x[110]+ x[111]+ x[112]+ x[113]+ x[130])/11611 - 1 ))^2)*1) +
    ((max(0,(1 - (x[77]+ x[85]+ x[108]+ x[109]+ x[110]+ x[111]+ x[112]+ x[113]+ x[130]+x[99])/11611 ))^2)*1) +
    ((max(0,((x[86])/12285 - 1 ))^2)*1) +
    ((max(0,(1 - (x[86]+x[87]+ x[91]+ x[94]+ x[97]+ x[100])/12285 ))^2)*1) +
    ((max(0,((x[102]+ x[118]+ x[119]+ x[121])/2712 - 1 ))^2)*1) +
    ((max(0,(1 - (x[102]+ x[118]+ x[119]+ x[121]+x[88]+ x[92]+ x[95]+ x[98]+ x[101]+ x[129])/2712 ))^2)*1) +
    ((max(0,((x[103]+x[120]+x[122])/635 - 1 ))^2)*1) +
    ((max(0,(1 - (x[103]+x[120]+x[122]+x[106]+x[107])/635 ))^2)*1) +
    ((max(0,((x[83]+ x[126])/1152 - 1 ))^2)*1) +
    ((max(0,(1 - (x[83]+ x[126]+x[80])/1152 ))^2)*1) +
    ((max(0,((x[84]+ x[87]+ x[91]+ x[94]+ x[96]+ x[97])/3374 - 1 ))^2)*1) +
    ((max(0,(1 - (x[84]+ x[87]+ x[91]+ x[94]+ x[96]+ x[97]+x[81])/3374 ))^2)*1) +
    ((max(0,((x[88]+ x[89]+ x[92]+ x[95]+ x[98]+ x[102]+ x[127])/2989 - 1 ))^2)*1) +
    ((max(0,(1 - (x[88]+ x[89]+ x[92]+ x[95]+ x[98]+ x[102]+ x[127]+x[82])/2989 ))^2)*1) +
    ((max(0,((x[90]+x[103])/1767 - 1 ))^2)*1)+
    #Bjerke
    ((max(0,((x[74]+x[76]+x[78]+x[85]+x[108]+x[75]+x[79]+x[86]+x[118]+x[119]+x[123]+x[120])/8088 - 1))^2)*1) +
    ((max(0,(1 - (x[74]+x[76]+x[78]+x[85]+x[108]+x[75]+x[79]+x[86]+x[118]+x[119]+x[123]+x[120]+x[83]+x[112]+x[84]+x[131]+x[114]+x[132]+x[115])/8088))^2)*1) +
    ((max(0,((x[109]+ x[111]+ x[126]+ x[128]+x[104]+ x[121]+ x[124]+ x[127]+x[105]+ x[122]+ x[125])/7188 - 1))^2)*1) +
    ((max(0,(1 - (x[109]+x[111]+x[126]+x[128]+x[104]+x[121]+x[124]+x[127]+x[105]+x[122]+x[125]+x[99]+x[108]+x[130]+x[100]+x[114]+x[101]+x[115])/7188))^2)*1)
}
Problem$lb <- rep(10, times = 64)
Problem$ub <- rep(1000000, times = 64)

outputfcn <- function(it, gbest, fx, x){
  cat(sprintf("Iteration: %d", it), sprintf("Error: %f", fx), "\n", sep = "\t");
  # return a negative value for PSwarm to abort execution
  1.0;
}


# Define a list with the solver options
Options <- list(
  cognitial = 0.5, fweight = 0.4, iweight = 0.9,
  maxf = 2000000000, maxit = 2000000000, size = 65 * Problem$Variables, iprint = 10000, social = 0.5,
  tol= 1E-6, ddelta = 0.5, idelta = 2,
  outputfcn = outputfcn, vectorized=0)

# Load the solver
#dyn.load("pswarm_r.so")  # Linux
dyn.load("pswarm_r")    # Windows

##########################
# Call the solver
Result <- .Call("pswarm_r", Problem, Options, .GlobalEnv)

# Results presents the obtained solution, its objective function value
# and a zero return value on success
print(Result)
View(Result[["x"]])
###########################
###########################
###########################
report <- data.frame(matrix(ncol = 66, nrow = 0))
for (i in 1:200){  
  # Call the solver
  set.seed(rnorm(1)*1E3)
  Result <- .Call("pswarm_r", Problem, Options, .GlobalEnv)
  cat("\n\nError=", Result$f, "\n\n")
  # print(Result)
  
  report<-rbind(report, append(Result$x, Result$f, after=0))
}

write.table(report, file = "./CA1_062021_1s10s_rest.csv", fileEncoding = "UTF-16", na = "", sep = ",", row.names = F,
            col.names = c("error", "x69",	"x70",	"x71",	"x72",	"x73",	"x74",	"x75",	"x76",	"x77",	"x78",	"x79",	"x80",	"x81",	"x82",	"x83",	"x84",	"x85",	"x86",	"x87",	"x88",	"x89",	"x90",	"x91",	"x92",	"x93",	"x94",	"x95",	"x96",	"x97",	"x98",	"x99",	"x100",	"x101",	"x102",	"x103",	"x104",	"x105",	"x106",	"x107",	"x108",	"x109",	"x110",	"x111",	"x112",	"x113",	"x114",	"x115",	"x116",	"x117",	"x118",	"x119",	"x120",	"x121",	"x122",	"x123",	"x124",	"x125",	"x126",	"x127",	"x128",	"x129",	"x130",	"x131",	"x132"))

rm(i, Options, Problem, outputfcn, Result)
#mid sep: change cossart2000, tyan, hajos, add enk eq, add coulin, ero-no x2&x5, change bezaire wts to 1, add jinnokosaka
#9/29: coulin & FT-enk eq are playing against each other; add or-lm to som+ equation/inequality; next solve IS eqs
#10/07: corrected ero numbers, added poolos assumption
#10/13: changed assumption to say 5:200 (5-10 plcs and 100s of pcs)
#12/02/20: changed biomarker wts to 1 after correcting a couple of volume errors. Modeling/Bezaire wts are at 5.
```
