## SupplementaryData for "Quantification of neuron types in the rodent hippocampal formation by data mining and numerical optimization": CA2_3.R

```
# Define a list with the problem definition
rm(Problem, outputfcn, Options, Result)
Problem <- list()
Problem$Variables <- 46
Problem$objf <- function(x) {
  ##################################
    (((x[24]/(x[23]*0.24) - 1)^2)*10) +
    #Ero
    ((x[40]/2143 - 1)^2) +
    #
    ((x[68]/1372 - 1)^2) +
    #
    (((x[25]+x[27])/67267 - 1)^2) +
    (((x[38]+x[46]+x[54]+x[55]+x[56])/23640 - 1)^2) +
    ((x[26]/57721 - 1)^2) +
    (((x[37]+x[39]+x[41]+x[42]+x[43]+x[44]+x[45]+x[49]+x[53])/863 - 1)^2) +
    (((x[23]+x[24])/697391 - 1)^2) +
    (((x[28]+x[30]+x[31]+x[33]+x[36]+x[48]+x[52])/10662 - 1)^2) +
    (((x[29]+x[32]+x[34]+x[35]+x[47]+x[50]+x[51]+x[57]+x[58])/10548 - 1)^2) +
    #
    (((x[40]/3844 - 1)^2)*10) +
    ((((x[25]+x[27]+x[38]+x[46]+x[54]+x[55]+x[56])/93015 - 1)^2)*10) +
    ((((x[26]+x[37]+x[39]+x[41]+x[42]+x[43]+x[44]+x[45]+x[49]+x[53])/45416 - 1)^2)*10) +
    ((((x[23]+x[24]+x[28]+x[30]+x[31]+x[33]+x[36]+x[48]+x[52])/254942 - 1)^2)*10) +
    ((((x[29]+x[32]+x[34]+x[35]+x[47]+x[50]+x[51]+x[57]+x[58])/128388 - 1)^2)*10) +
    (((x[40]/2246 - 1)^2)*10) +
    ((((x[25]+x[27]+x[38]+x[46]+x[54]+x[55]+x[56])/34220 - 1)^2)*10) +
    ((((x[26]+x[37]+x[39]+x[41]+x[42]+x[43]+x[44]+x[45]+x[49]+x[53])/21803 - 1)^2)*10) +
    ((((x[29]+x[32]+x[34]+x[35]+x[47]+x[50]+x[51]+x[57]+x[58])/72398 - 1)^2)*10) +
    (((x[23]+x[24]+x[59])/667800 - 1)^2) +
    ((((x[23]+x[24]+x[28]+x[30]+x[31]+x[33]+x[36]+x[48]+x[52]+x[59]+x[60]+x[61]+x[62]+x[63])/360000 - 1)^2)*10) +
    ((((x[23]+x[24]+x[28]+x[30]+x[31]+x[33]+x[36]+x[48]+x[52]+x[59]+x[60]+x[61]+x[62]+x[63])/360000 - 1)^2)*10) +
    ((((x[23]+x[24]+x[28]+x[30]+x[31]+x[33]+x[36]+x[48]+x[52]+x[59]+x[60]+x[61]+x[62]+x[63])/250400 - 1)^2)*10) +
    ((((x[23]+x[24]+x[28]+x[30]+x[31]+x[33]+x[36]+x[48]+x[52]+x[59]+x[60]+x[61]+x[62]+x[63])/225000 - 1)^2)*10) +
    ((((x[23]+x[24]+x[28]+x[30]+x[31]+x[33]+x[36]+x[48]+x[52]+x[59]+x[60]+x[61]+x[62]+x[63])/188000 - 1)^2)*10) +
    ((((x[23]+x[24]+x[28]+x[30]+x[31]+x[33]+x[36]+x[48]+x[52]+x[59]+x[60]+x[61]+x[62]+x[63])/414800 - 1)^2)*10) +
    ((((x[23]+x[24]+x[28]+x[30]+x[31]+x[33]+x[36]+x[48]+x[52]+x[59]+x[60]+x[61]+x[62]+x[63])/181440 - 1)^2)*10) +
    ((((x[23]+x[24]+x[28]+x[30]+x[31]+x[33]+x[36]+x[48]+x[52]+x[59]+x[60]+x[61]+x[62]+x[63])/416498 - 1)^2)*10) +
    (((x[23]+x[24]+x[25]+x[26]+x[27]+x[28]+x[29]+x[30]+x[31]+x[32]+x[33]+x[34]+x[35]+x[36]+x[37]+x[38]+x[39]+x[40]+x[41]+x[42]+x[43]+x[44]+x[45]+x[46]+x[47]+x[48]+x[49]+x[50]+x[51]+x[52]+x[53]+x[54]+x[55]+x[56]+x[57]+x[58])/876739 - 1)^2) +
    (((2*(x[23]+x[24]))/(x[23]+x[24]+x[28]+x[30]+x[43]+x[47]+x[49]) - 1)^2) +
    (((55*(x[30]))/(15*(x[28]+x[30]+x[43]+x[47]+x[49])) - 1)^2) +
    (((55*x[28])/(18*(x[28]+x[30]+x[43]+x[47]+x[49])) - 1)^2) +
    (((55*(x[47]+x[49]))/(14*(x[28]+x[30]+x[43]+x[47]+x[49])) - 1)^2) +
    (((55*x[43])/(8*(x[28]+x[30]+x[43]+x[47]+x[49])) - 1)^2) +
    (((135*(x[31]+x[41]))/(38*(x[31]+x[33]+x[41]+x[45]+x[46]+x[49]+x[55]+x[56])) - 1)^2) +
    (((135*(x[45]+x[46]+x[49]))/(32*(x[31]+x[33]+x[41]+x[45]+x[46]+x[49]+x[55]+x[56])) - 1)^2) +
    (((135*x[55])/(52*(x[31]+x[33]+x[41]+x[45]+x[46]+x[49]+x[55]+x[56])) - 1)^2) +
    (((135*x[56])/(11*(x[31]+x[33]+x[41]+x[45]+x[46]+x[49]+x[55]+x[56])) - 1)^2) +
    (((135*(x[33]))/(2*(x[31]+x[33]+x[41]+x[45]+x[46]+x[49]+x[55]+x[56])) - 1)^2) +
    (((8*(x[47]+x[48]+x[49]))/(x[45]+x[46]) - 1)^2) +
    #Miles
    #(((8*(x[28]+x[29]+x[30]+x[31]))/(17*(x[32]+x[33])) - 1)^2) +
    (((8*(x[28]+x[30]+x[31]))/(17*(x[32]+x[33])) - 1)^2) +
    #Gulyas
    (((30*(x[28]))/(31*(x[30])) - 1)^2) +
    (((25*(x[30]))/(13*(x[28]+x[30]+x[31])) - 1)^2) +
    (((25*(x[28]))/(12*(x[28]+x[30]+x[31])) - 1)^2) +
    (((5*x[31])/(2*(x[28]+x[30]+x[31])) - 1)^2) +
    #
    #(((30*(x[28]+x[29]))/(31*(x[30])) - 1)^2) +
    #(((25*(x[30]))/(13*(x[28]+x[29]+x[30]+x[31])) - 1)^2) +
    #(((25*(x[28]+x[29]))/(12*(x[28]+x[29]+x[30]+x[31])) - 1)^2) +
    #(((5*x[31])/(2*(x[28]+x[29]+x[30]+x[31])) - 1)^2) +
    #
    (((758*(x[26]+x[27]))/(x[23]+x[24]) - 1)^2) +
    (((15*x[44])/(11*(x[39]+x[41]+x[44])) - 1)^2) +
    (((15*x[41])/(2*(x[39]+x[41]+x[44])) - 1)^2) +
    (((15*x[39])/(2*(x[39]+x[41]+x[44])) - 1)^2) +
    (((9*(x[45]+x[46]))/(5*(x[39]+x[45]+x[46])) - 1)^2) +
    (((9*x[39])/(4*(x[39]+x[45]+x[46])) - 1)^2) +
    (((115*(x[30]))/(12*(x[30]+x[31]+x[39]+x[44]+x[45]+x[46])) - 1)^2) +
    (((115*x[31])/(24*(x[30]+x[31]+x[39]+x[44]+x[45]+x[46])) - 1)^2) +
    (((115*x[39])/(19*(x[30]+x[31]+x[39]+x[44]+x[45]+x[46])) - 1)^2) +
    (((115*x[44])/(37*(x[30]+x[31]+x[39]+x[44]+x[45]+x[46])) - 1)^2) +
    (((115*(x[45]+x[46]))/(23*(x[30]+x[31]+x[39]+x[44]+x[45]+x[46])) - 1)^2) +
    (((10*x[60])/(3*(x[60]+x[61]+x[62])) - 1)^2) +
    (((2*x[61])/(x[60]+x[61]+x[62]) -1)^2) +
    (((10*x[62])/(2*(x[60]+x[61]+x[62])) - 1)^2) +
    (((9*x[60])/(7*x[61]) - 1)^2) +
    (((4*x[62])/(6*x[63]) - 1)^2) +
    #Ero
    ((x[59]/66751 - 1)^2) +
    (((x[60]+x[61]+x[62]+x[63])/291 - 1)^2) +
    #
    ((x[64]/2327 - 1)^2) +
    ((x[65]/4296 - 1)^2) +
    ((x[66]/923 - 1)^2) +
    ((x[67]/2567 - 1)^2) +
    ((((x[59]+x[60]+x[61]+x[62]+x[63])/19221 - 1)^2)*10) +
    #Kosaka
    (map(0,(x[29]/(0.26*(x[29]+x[32]+x[34]+x[35]+x[47]+x[50]+x[51]+x[57]+x[58])) - 1)^2)*1) +
    (map(0,(1 - (x[29]+x[32]+x[34]+x[47]+x[50]+x[58])/(0.26*(x[29]+x[32]+x[34]+x[35]+x[47]+x[50]+x[51]+x[57]+x[58])))^2)*1) +
    (map(0,((x[28]+x[30]+ x[60]+ x[61]+ x[62])/(0.45*(x[28]+x[30]+x[31]+x[33]+x[36]+x[48]+x[52])) - 1)^2)*1) +
    (map(0,(1 - (x[28]+x[30]+x[60]+x[61]+x[62]+x[33]+x[48])/(0.45*(x[28]+x[30]+x[31]+x[33]+x[36]+x[48]+x[52])))^2)*1) +
    (map(0,(1 - (x[41]+x[42]+x[43]+x[49])/(0.18*(x[37]+x[39]+x[41]+x[42]+x[43]+x[44]+x[45]+x[49]+x[53])))^2)*1) +
    #Jinno & Kim
    ((map(0,(1- (x[29]+x[32]+x[34]+x[47]+x[50]+x[58])/2951))^2)*1) +
    ((map(0,(x[29]/2951 - 1))^2)*1) +
    ((map(0,(1 - (x[28]+x[30]+x[60]+x[61]+x[62]+x[33]+x[48])/6021))^2)*1) +
    ((map(0,((x[28]+x[30]+x[60]+x[61]+x[62])/6021 - 1))^2)*1) +
    ((map(0,(1 - (x[41]+x[42]+x[43]+x[49])/272 ))^2)*1) +
    ((map(0,(1 - (x[29]+x[32]+x[34]+x[47]+x[50]+x[58])/3684))^2)*1) +
    ((map(0,(x[29]/3684 -1))^2)*1) +
    ((map(0,(1 - (x[28]+x[30]+x[33]+x[48])/2974))^2)*1) +
    ((map(0,((x[28]+x[30])/2974 -1))^2)*1) +
    ((map(0,((x[60]+x[61]+x[62])/472 -1))^2)*1) +
    ((map(0,(1 - (x[41]+x[42]+x[43]+x[49])/411))^2)*1) +
    ((map(0,(1 - (x[32]+x[34]+x[35]+x[47]+x[50]+x[58])/313))^2)*1) +
    ((map(0,(1 - (x[33]+x[36]+x[48]+x[63])/1600))^2)*1) +
    ((map(0,(1 - (x[37]+x[39]+x[41]+x[42]+x[43]+x[49])/147))^2)*1) +
    ((map(0,(1 - (x[38]+x[56])/957))^2)*1) +
    ((map(0,(1 - x[40]/16))^2)*1) +
    ((map(0,(1 - (x[57]+x[32]+x[34]+x[47]+x[50]+x[58])/3191))^2)*1) +
    ((map(0,(x[57]/3191 -1))^2)*1) +
    ((map(0,(1 - (x[33]+x[48]+x[63])/2521))^2)*1) +
    ((map(0,(1 - (x[41]+x[42]+x[43]+x[49])/209))^2)*1) +
    ((map(0,(1 - (x[55]+x[56])/3839))^2)*1) +
    ((map(0,(x[55]/3839 -1))^2)*1) +
    ((map(0,(1 - x[40]/218))^2)*1) +
    ((map(0,(1 - (x[29]+x[32]+x[34]+x[35]+x[47]+x[50]+x[58])/5120))^2)*1) +
    ((map(0,(1 - (x[30]+x[31]+x[33]+x[36]+x[48]+x[60]+x[61]+x[62]+x[63])/8590))^2)*1) +
    ((map(0,(1 - (x[39]+x[44]+x[37]+x[41]+x[42]+x[43]+x[45]+x[49])/915))^2)*1) +
    ((map(0,((x[39]+x[44])/915 -1))^2)*1) +
    ((map(0,(1 - (x[55]+x[38]+x[46]+x[56])/4298))^2)*1) +
    ((map(0,(x[55]/4298 -1))^2)*1) +
    ((map(0,(1 - x[40]/24))^2)*1) +
    ((map(0,(1 - (x[29]+x[32]+x[34]+x[35]+x[47]+x[50]+x[57]+x[58])/3816))^2)*1) +
    ((map(0,(1 - (x[30]+x[33]+x[36]+x[48]+x[60]+x[61]+x[62]+x[63])/3558))^2)*1) +
    ((map(0,(1 - (x[44]+x[37]+x[41]+x[42]+x[43]+x[49])/367))^2)*1) +
    ((map(0,(x[44]/367 -1))^2)*1) +
    ((map(0,(1 - (x[38]+x[56])/1407))^2)*1) +
    ((map(0,(1 - x[40]/1))^2)*1) +
    ((map(0,(1 - (x[29]+x[32]+x[34]+x[35]+x[47]+x[50]+x[57]+x[58])/9550))^2)*1) +
    ((map(0,(1 - (x[30]+x[33]+x[36]+x[48])/10906))^2)*1) +
    ((map(0,(1 - (x[60]+x[61]+x[62]+x[63])/413))^2)*1) +
    ((map(0,(1 - (x[44]+x[37]+x[41]+x[42]+x[43]+x[49])/1818))^2)*1) +
    ((map(0,(x[44]/1818 -1))^2)*1) +
    ((map(0,(1 - (x[38]+x[56])/4562))^2)*1) +
    ((map(0,(1 - x[40]/33))^2)*1) +
    ((map(0,(1 - (x[51]+x[32]+x[34]+x[35]+x[47]+x[50]+x[58])/1011))^2)*1) +
    ((map(0,(x[51]/1011 -1))^2)*1) +
    ((map(0,(1 - (x[31]+x[52]+x[33]+x[36]+x[48])/1348))^2)*1) +
    ((map(0,((x[31]+x[52])/1348 -1))^2)*1) +
    ((map(0,(1 - (x[45]+x[53]+x[37]+x[41]+x[42]+x[43]+x[49])/362))^2)*1) +
    ((map(0,((x[45]+x[53])/362 -1))^2)*1) +
    ((map(0,(1 - (x[46]+x[54]+x[38]+x[56])/2125))^2)*1) +
    ((map(0,((x[46]+x[54])/2125 -1))^2)*1) +
    ((map(0,(1 - x[40]/5))^2)*1) +
    ((map(0,(1 - (x[35]+x[32]+x[34]+x[47]+x[50]+x[58])/146))^2)*1) +
    ((map(0,(x[35]/146 -1))^2)*1) +
    ((map(0,(1 - (x[36]+x[33]+x[48]+x[63])/1105))^2)*1) +
    ((map(0,(x[36]/1105 -1))^2)*1) +
    ((map(0,(1 - (x[37]+x[39]+x[41]+x[42]+x[43]+x[44]+x[45]+x[49])/160))^2)*1) +
    ((map(0,(x[37]/160 -1))^2)*1) +
    ((map(0,(1 - (x[38]+x[46]+x[56])/708))^2)*1) +
    ((map(0,(x[38]/708 -1))^2)*1) +
    ((map(0,(1 - x[40]/12))^2)*1) +
    ((map(0,(1 - (x[35]+x[32]+x[34]+x[47]+x[50]+x[58])/614))^2)*1) +
    ((map(0,(x[35]/614 -1))^2)*1) +
    ((map(0,(1 - (x[36]+x[33]+x[48])/987))^2)*1) +
    ((map(0,(1 - (x[63])/194))^2)*1) +
    ((map(0,(x[36]/987 -1))^2)*1) +
    ((map(0,(1 - (x[37]+x[39]+x[41]+x[42]+x[43]+x[44]+x[45]+x[49])/233))^2)*1) +
    ((map(0,(x[37]/233 -1))^2)*1) +
    ((map(0,(1 - (x[38]+x[46]+x[56])/750))^2)*1) +
    ((map(0,(x[38]/750 -1))^2)*1) +
    ((map(0,(1 - x[40]/13))^2)*1) +
    ((map(0,(1 - (x[29]+x[32]+x[34]+x[35]+x[47]+x[50]+x[51]+x[57]+x[58])/949))^2)*1) +
    ((map(0,(1 - (x[28]+x[30]+x[31]+x[33]+x[36]+x[48]+x[52]+x[60]+x[61]+x[62]+x[63])/4877))^2)*1) +
    ((map(0,(1 - (x[37]+x[39]+x[41]+x[42]+x[43]+x[44]+x[45]+x[49]+x[53])/334))^2)*1) +
    ((map(0,(1 - (x[55]+x[38]+x[46]+x[54]+x[56])/2728))^2)*1) +
    ((map(0,(x[55]/2728 -1))^2)*1) +
    ((map(0,(1 - x[40]/367))^2)*1)+
    ((map(0,(1- x[66]/388))^2)*1)+
    ((map(0,(1- x[67]/144))^2)*1)+
    ((map(0,(1- x[65]/17))^2)*1)+
    ((map(0,(1- x[66]/699))^2)*1)+
    ((map(0,(1- x[67]/205))^2)*1)+
    ((map(0,(1- x[65]/55))^2)*1)+
    ((map(0,(1- x[66]/75))^2)*1)+
    ((map(0,(1- x[67]/138))^2)*1)+
    ((map(0,(1- x[65]/50))^2)*1)+
    ((map(0,((x[28]+x[29]+x[30])/4678 - 1))^2)*1)+
    ((map(0,(1 - (x[29]+x[32]+x[34]+x[47]+x[50]+x[58]+x[28]+x[30]+x[33]+x[48]+x[41]+x[42]+x[43]+x[49])/4678))^2)*1)+
    ((map(0,((x[55]+x[57])/1998 - 1))^2)*1)+
    ((map(0,(1 - (x[32]+x[34]+x[47]+x[50]+x[58]+x[33]+x[48]+x[63]+x[41]+x[42]+x[43]+x[49]+x[56]+x[40]+x[55]+x[57])/1998))^2)*1)+
    ((map(0,((x[60]+x[61]+x[62])/478 - 1))^2)*1)+
    ((map(0,(1 - (x[66]+x[60]+x[61]+x[62]+x[67]+x[65])/478))^2)*1)+
    ((map(0,(1 - x[63]/102))^2)*1)
}

Problem$lb <- rep(10, times = 46)
Problem$ub <- rep(1000000, times = 46)

outputfcn <- function(it, gbest, fx[, x[){
  cat(sprintf("Iteration: %d", it), sprintf("Error: %f", fx[), "\n", sep = "\t");
  # return a negative value for PSwarm to abort ex[ecution
  1.0;
}


# Define a list with the solver options
Options <- list(
  cognitial = 0.5, fweight = 0.4, iweight = 0.9,
  max[f = 2000000000, max[it = 2000000000, size = 47 * Problem$Variables, iprint = 10000, social = 0.5,
  tol= 1E-20, ddelta = 0.5, idelta = 2,
  outputfcn = outputfcn, vectorized=0)

# Load the solver
#dyn.load("pswarm_r.so")  # Linux[
dyn.load("pswarm_r")    # Windows

##########################
# Call the solver
Result <- .Call("pswarm_r", Problem, Options, .GlobalEnv)

# Results presents the obtained solution, its objective function value
# and a zero return value on success
print(Result)

###########################
report <- data.frame(matrix[(ncol = 47, nrow = 0))
for (i in 1:100){  
  # Call the solver
  set.seed(rnorm(1)*1E3)
  Result <- .Call("pswarm_r", Problem, Options, .GlobalEnv)
  cat("\n\nError=", Result$f, "\n\n")
  # print(Result)
  
  report<-rbind(report, append(Result$x[, Result$f, after=0))
}

write.table(report, file = "./CA32_021921_2.csv", fileEncoding = "UTF-16", na = "", sep = ",", row.names = F,
            col.names = c("error", "x[23]","x[24]","x[25]","x[26]","x[27]","x[28]","x[29]","x[30]","x[31]","x[32]","x[33]","x[34]","x[35]","x[36]","x[37]","x[38]","x[39]","x[40]","x[41]","x[42]","x[43]","x[44]","x[45]","x[46]","x[47]","x[48]","x[49]","x[50]","x[51]","x[52]","x[53]","x[54]","x[55]","x[56]","x[57]","x[58]","x[59]","x[60]","x[61]","x[62]","x[63]","x[64]","x[65]","x[66]","x[67]","x[68]"))

rm(i, Options, Problem, outputfcn, Result)
```
