## SupplementaryData for "Quantification of neuron types in the rodent hippocampal formation by data mining and numerical optimization": DG.r

```
# Define a list with the problem definition
rm(Problem, outputfcn, Options, Result)
Problem <- list()
Problem$Variables <- 22

Problem$objf <- function(x) {
  ##################################
  ((((x[19]+x[20])/(2*x[10] + 2*x[16] + 2*x[18]) - 1 )^2)*1) +
    ((((11*x[19])/(6*x[20]) - 1 )^2)*1) +
    ((((6*x[4])/(35*x[5]) - 1 )^2)*1) +
    (((x[4]/(3*x[5]) - 1 )^2)*1) +
    ((((17*x[19])/(14*x[20]) - 1 )^2)*1) +
    ((((x[8]+x[9])/(6*x[7]) - 1 )^2)*1) +
    ((((5*x[7])/(x[6]+x[11]+x[12]+x[13]+x[14]+x[15]+x[17]+x[21]+x[22]) - 1 )^2)*1) +
    ((((7*(x[8]+x[9]))/(6*(x[13]+x[15])) - 1 )^2)*1) +
    ((((4*(x[13]+x[15]))/(7*(x[6]+x[11])) - 1 )^2)*1) +
    ((((2*(x[8]+x[9]))/(x[6]+x[11]+x[12]) - 1 )^2)*1) +
    ((((3*(x[6]+x[11]+x[12]))/(4*(x[13]+x[14]+x[21])) - 1 )^2)*1) +
    ((((11*(x[13]+x[14]+x[21]))/(6*x[17]) - 1 )^2)*1) +
    ((((x[17])/(x[22]+x[15]) - 1 )^2)*1) +
    ((((64*(x[16]+x[18]))/(169*x[3]) - 1 )^2)*1) +
    ((((x[6]/16801) + (x[11]/16801) + (x[13]/16801) + (x[15]/16801) - 1 )^2)*10) +
    ((((x[2]/46734) + (x[4]/46734) + (x[5]/46734) + (x[6]/46734) + (x[11]/46734) + (x[13]/46734) + (x[15]/46734) - 1 )^2)*10) +
    ((((x[6]/35900) + (x[7]/35900) + (x[8]/35900) + (x[9]/35900) + (x[10]/35900) + (x[11]/35900) + (x[12]/35900) + (x[13]/35900) + (x[14]/35900) + (x[15]/35900) + (x[16]/35900) + (x[17]/35900) + (x[18]/35900) + (x[19]/35900) + (x[20]/35900) + (x[21]/35900) + (x[22]/35900) - 1 )^2)*1) +
    #121420: 
    ((((47*(x[7]+x[8]+x[9]+x[12]+x[14]+x[17]+x[21]+x[22]))/(26*(x[6]+x[11]+x[13]+x[15])) - 1 )^2)*1) +
    #121420: 
    ((((27*(x[6]+x[11]+x[13]+x[15]))/(47*(x[10]+x[16]+x[18]+x[19]+x[20])) - 1 )^2)*1) +
    ((((x[2]/90) - 1 )^2)*10) +
    ((((x[4]/21542) + (x[5]/21542) - 1 )^2)*10) +
    ((((x[6]/16251) + (x[11]/16251) + (x[13]/16251) + (x[15]/16251) - 1 )^2)*10) +
    ((((x[1]/1200000) + (x[7]/1200000) + (x[8]/1200000) + (x[9]/1200000) + (x[12]/1200000) + (x[14]/1200000) + (x[17]/1200000) + (x[21]/1200000) + (x[22]/1200000) - 1 )^2)*10) +
    ((((x[1])/(49*(x[7]+x[8]+x[9]+x[12]+x[14]+x[17]+x[21]+x[22])) - 1 )^2)*10) +
    #((((42*x[3])/(58*(x[10]+x[16]+x[18]+x[19]+x[20])) - 1 )^2)*1) +
    #((((42*(x[1]+x[2]+x[3]+x[4]+x[5]))/(58*(x[6]+x[7]+x[8]+x[9]+x[10]+x[11]+x[12]+x[13]+x[14]+x[15]+x[16]+x[17]+x[18]+x[19]+x[20]+x[21]+x[22])) - 1 )^2)*1) +
    #((((x[1]/1258848) + (x[2]/1258848) + (x[3]/1258848) + (x[4]/1258848) + (x[5]/1258848) + (x[6]/1258848) + (x[7]/1258848) + (x[8]/1258848) + (x[9]/1258848) + (x[10]/1258848) + (x[11]/1258848) + (x[12]/1258848) + (x[13]/1258848) + (x[14]/1258848) + (x[15]/1258848) + (x[16]/1258848) + (x[17]/1258848) + (x[18]/1258848) + (x[19]/1258848) + (x[20]/1258848) + (x[21]/1258848) + (x[22]/1258848) - 1 )^2)*1) +
    ((((x[3]/147050) + (x[10]/147050) + (x[16]/147050) +(x[18]/147050) + (x[19]/147050) + (x[20]/147050) - 1 )^2)*10) +
    ((((x[2]/44295) + (x[4]/44295) + (x[5]/44295) + (x[6]/44295) + (x[11]/44295) + (x[13]/44295) + (x[15]/44295) - 1 )^2)*10) +
    #ero granule cells
    ((((x[1]/747485) - 1 )^2)*1) +
    ((((x[1]/1080000) + (x[7]/1080000) + (x[8]/1080000) + (x[9]/1080000) + (x[12]/1080000) + (x[14]/1080000) + (x[17]/1080000) + (x[21]/1080000) + (x[22]/1080000) - 1 )^2)*10) +
    ((((x[1]/1065000) + (x[7]/1065000) + (x[8]/1065000) + (x[9]/1065000) + (x[12]/1065000) + (x[14]/1065000) + (x[17]/1065000) + (x[21]/1065000) + (x[22]/1065000) - 1 )^2)*10) +
    ((((x[1]/1180000) + (x[7]/1180000) + (x[8]/1180000) + (x[9]/1180000) + (x[12]/1180000) + (x[14]/1180000) + (x[17]/1180000) + (x[21]/1180000) + (x[22]/1180000) - 1 )^2)*10) +
    ((((x[1]/969900) + (x[7]/969900) + (x[8]/969900) + (x[9]/969900) + (x[12]/969900) + (x[14]/969900) + (x[17]/969900) + (x[21]/969900) + (x[22]/969900) - 1 )^2)*10) +
    ((((x[1]/1172290) + (x[7]/1172290) + (x[8]/1172290) + (x[9]/1172290) + (x[12]/1172290) + (x[14]/1172290) + (x[17]/1172290) + (x[21]/1172290) + (x[22]/1172290) - 1 )^2)*10) +
    ((((x[1]/1200000) + (x[7]/1200000) + (x[8]/1200000) + (x[9]/1200000) + (x[12]/1200000) + (x[14]/1200000) + (x[17]/1200000) + (x[21]/1200000) + (x[22]/1200000) - 1 )^2)*10) +
    ((((x[1]/1756098) + (x[7]/1756098) + (x[8]/1756098) + (x[9]/1756098) + (x[12]/1756098) + (x[14]/1756098) + (x[17]/1756098) + (x[21]/1756098) + (x[22]/1756098) - 1 )^2)*10) +
    ((((x[1]/1073171) + (x[7]/1073171) + (x[8]/1073171) + (x[9]/1073171) + (x[12]/1073171) + (x[14]/1073171) + (x[17]/1073171) + (x[21]/1073171) + (x[22]/1073171) - 1 )^2)*10) +
    #ero gl ins
    ((((x[7]/13832) + (x[8]/13832) + (x[9]/13832) + (x[12]/13832) + (x[14]/13832) + (x[17]/13832) + (x[21]/13832) + (x[22]/13832) - 1 )^2)*1) +
    ((((x[2]/65420) + (x[4]/65420) + (x[5]/65420) + (x[6]/65420) + (x[11]/65420) + (x[13]/65420) + (x[15]/65420) - 1 )^2)*10) +
    ((((x[2]/52495) + (x[4]/52495) + (x[5]/52495) + (x[6]/52495) + (x[11]/52495) + (x[13]/52495) + (x[15]/52495) - 1 )^2)*10) +
    ((((x[2]/53000) + (x[4]/53000) + (x[5]/53000) + (x[6]/53000) + (x[11]/53000) + (x[13]/53000) + (x[15]/53000) - 1 )^2)*10) +
    ((((x[2]/63200) + (x[4]/63200) + (x[5]/63200) + (x[6]/63200) + (x[11]/63200) + (x[13]/63200) + (x[15]/63200) - 1 )^2)*10) +
    ((((x[2]/49275) + (x[4]/49275) + (x[5]/49275) + (x[6]/49275) + (x[11]/49275) + (x[13]/49275) + (x[15]/49275) - 1 )^2)*10) +
    ((((x[2]/45000) + (x[4]/45000) + (x[5]/45000) + (x[6]/45000) + (x[11]/45000) + (x[13]/45000) + (x[15]/45000) - 1 )^2)*10) +
    ((((x[2]/40000) + (x[4]/40000) + (x[5]/40000) + (x[6]/40000) + (x[11]/40000) + (x[13]/40000) + (x[15]/40000) - 1 )^2)*10) +
    #ero hilar excitatory
    ((((x[2]/149234) + (x[4]/149234) + (x[5]/149234) - 1 )^2)*1) +
    #ero hilar ins
    ((((x[6]/4724) + (x[11]/4724) + (x[13]/4724) + (x[15]/4724) - 1)^2)*1) +
    #ero ml excitatory
    (((x[3]/302448)^2)*1) +
    #ero ml ins smi & smo
    #((((x[10]/12543) + (x[16]/12543) + (x[18]/12543) - 1)^2)*1) +
    #((((x[19]/25085) + (x[20]/25085) - 1)^2)*1) +
    ((((x[10]/37628) + (x[16]/37628) + (x[18]/37628) + (x[19]/37628) + (x[20]/37628) - 1)^2)*1) +
    ((((x[3]/378459) + (x[10]/378459) + (x[16]/378459) +(x[18]/378459) + (x[19]/378459) + (x[20]/378459) - 1 )^2)*10) +
    ((((x[2]/127550) + (x[4]/127550) + (x[5]/127550) + (x[6]/127550) + (x[11]/127550) + (x[13]/127550) + (x[15]/127550) - 1 )^2)*10) +
    ((((x[1]/648505) + (x[7]/648505) + (x[8]/648505) + (x[9]/648505) + (x[12]/648505) + (x[14]/648505) + (x[17]/648505) + (x[21]/648505) + (x[22]/648505) - 1 )^2)*10) +
    #harney
    #((((39*x[7])/(6*x[8]) - 1 )^2)*1) +
    #121420
    #((((34*(x[13]+x[14]))/(6*(x[6]+x[11]+x[12])) - 1 )^2)*1) +
    #
    ((((30*x[5])/(4*x[4]) - 1 )^2)*1) +
    ((((65*(x[7]+x[8]))/(13*(x[6]+x[7]+x[8]+x[11]+x[12]+x[13]+x[14]+x[15]+x[17]+x[18]+x[19]+x[20]+x[21]+x[22])) - 1 )^2)*1) +
    ((((65*(x[6]+x[11]+x[12]))/(11*(x[6]+x[7]+x[8]+x[11]+x[12]+x[13]+x[14]+x[15]+x[17]+x[18]+x[19]+x[20]+x[21]+x[22])) - 1 )^2)*1) +
    ((((65*(x[13]+x[14]+x[21]))/(10*(x[6]+x[7]+x[8]+x[11]+x[12]+x[13]+x[14]+x[15]+x[17]+x[18]+x[19]+x[20]+x[21]+x[22])) - 1 )^2)*1) +
    ((((65*(x[15]+x[17]+x[22]))/(17*(x[6]+x[7]+x[8]+x[11]+x[12]+x[13]+x[14]+x[15]+x[17]+x[18]+x[19]+x[20]+x[21]+x[22])) - 1 )^2)*1) +
    ((((65*(x[18]+x[19]+x[20]))/(14*(x[6]+x[7]+x[8]+x[11]+x[12]+x[13]+x[14]+x[15]+x[17]+x[18]+x[19]+x[20]+x[21]+x[22])) - 1 )^2)*1) +
    ((((39*(x[11]+x[12]))/(6*(x[6]+x[8]+x[11]+x[12]+x[13]+x[14]+x[17]+x[21])) - 1)^2)*1) +
    ((((39*x[6])/(14*(x[6]+x[8]+x[11]+x[12]+x[13]+x[14]+x[17]+x[21])) - 1 )^2)*1) +
    ((((39*(x[13]+x[14]))/(9*(x[6]+x[8]+x[11]+x[12]+x[13]+x[14]+x[17]+x[21])) - 1)^2)*1) +
    ((((39*(x[17]+x[21]))/(4*(x[6]+x[8]+x[11]+x[12]+x[13]+x[14]+x[17]+x[21])) - 1 )^2)*1) +
    ((((39*x[8])/(6*(x[6]+x[8]+x[11]+x[12]+x[13]+x[14]+x[17]+x[21])) - 1 )^2)*1) +
    #Soltez
    ((((x[1]/1000000) - 1)^2)*5) +
    ((((x[4]/30000) - 1)^2)*5) +
    #(((((x[8]+x[9]+x[10])/10000) - 1)^2)*5) +
    (((((x[8]+x[10])/10000) - 1)^2)*5) +
    (((((x[13]+x[14])/12000) - 1)^2)*5) +
    #new kosaka
    #((((x[6]+x[11]+x[13]+x[15])/(0.43*(x[2]+x[4]+x[5]+x[6]+x[11]+x[13]+x[15])) - 1)^2)*5) +
    #((((x[7]+x[8]+x[9]+x[12]+x[14]+x[17]+x[21]+x[22])/(0.55*(x[1]+x[7]+x[8]+x[9]+x[12]+x[14]+x[17]+x[21]+x[22])) - 1)^2)*5) +
    #((((x[10]+x[16]+x[18]+x[19]+x[20])/(0.03*(x[3]+x[10]+x[16]+x[18]+x[19]+x[20])) - 1)^2)*5) +
    #new kosaka
    ((max(0,(1 - ((x[6]+x[15])/(0.17*(x[6]+x[11]+x[13]+x[15])))))^2)*1)+
    ((max(0,(((x[7]+x[8])/(0.25*(x[7]+x[8]+x[9]+x[12]+x[14]+x[17]+x[21]+x[22]))) - 1))^2)*1)+
    ((max(0,(1 - ((x[7]+x[8]+x[17]+x[21]+x[22])/(0.25*(x[7]+x[8]+x[9]+x[12]+x[14]+x[17]+x[21]+x[22])))))^2)*1)+
    ((max(0,(1 - ((x[16]+x[20])/(0.03*(x[10]+x[16]+x[18]+x[19]+x[20])))))^2)*1)+
    #Jinno & Kim
    ((max(0,(1-((x[6]+x[15])/673)))^2)*1)+
    ((max(0,((x[7]+x[8])/3680 - 1))^2)*1)+
    ((max(0,(1-((x[7]+x[8]+x[17]+x[21]+x[22])/3680)))^2)*1)+
    ((max(0,(1- (x[16]+x[20])/541))^2)*1)+
    ((max(0,(1- (x[6]+x[15])/732))^2)*1)+
    ((max(0,((x[7]+x[8])/985 - 1))^2)*1)+
    ((max(0,(1-(x[7]+x[8]+x[17]+x[21]+x[22])/985))^2)*1)+
    ((max(0,(1- (x[16]+x[20])/393))^2)*1)+
    ((max(0,((x[13]/4963) - 1))^2)*1)+
    ((max(0,(1-(x[13]+x[6]+x[11]+x[15])/4963))^2)*1)+
    ((max(0,(x[14]/139 - 1))^2)*1)+
    ((max(0,(1-(x[12]+x[14])/139))^2)*1)+
    ((max(0,(x[13]/7166 - 1))^2)*1)+
    ((max(0,(1-(x[13]+x[6]+x[11]+x[15])/7166))^2)*1)+
    ((max(0,(x[14]/3954 - 1))^2)*1)+
    ((max(0,(1-(x[12]+x[14])/3954))^2)*1)+
    ((max(0,(1-(x[6]+x[15])/763))^2)*1)+
    ((max(0,(1-(x[9]+x[17]+x[21]+x[22])/1317))^2)*1)+
    ((max(0,(1-(x[10]+x[16]+x[18]+x[19]+x[20])/198))^2)*1)+
    ((max(0,(1-(x[6]+x[15])/993 - 1))^2)*1)+
    ((max(0,(1-(x[9]+x[17]+x[21]+x[22])/1066))^2)*1)+
    ((max(0,(1-(x[10]+x[16]+x[18]+x[19]+x[20])/666 ))^2)*1)+
    ((max(0,(1-(x[17]+x[21]+x[22])/1605 ))^2)*1)+
    ((max(0,(1- x[16]/368))^2)*1)+
    ((max(0,(1-(x[6]+x[15])/559 ))^2)*1)+
    ((max(0,(1-(x[17]+x[21]+x[22])/322 ))^2)*1)+
    ((max(0,(1- x[16]/542))^2)*1)+
    ((max(0,((x[13]/5956) - 1))^2)*1)+
    ((max(0,(1-(x[6]+x[11]+x[15]+x[13])/5956))^2)*1)+
    ((max(0,((x[14]/1544) - 1))^2)*1)+
    ((max(0,(1-(x[7]+x[8]+x[9]+x[12]+x[17]+x[21]+x[22]+x[14])/1544))^2)*1)+
    ((max(0,((x[18]+x[19]+x[20])/810 - 1))^2)*1)+
    ((max(0,(1-(x[10]+x[16]+x[18]+x[19]+x[20])/810))^2)*1)+
    ((max(0,(x[11]/149 - 1))^2)*1)+
    ((max(0,(1-(x[6]+x[13]+x[15]+x[11])/149))^2)*1)+
    ((max(0,((x[9]+x[12])/541 - 1))^2)*1)+
    ((max(0,(1-(x[14]+x[17]+x[21]+x[22]+x[9]+x[12])/541))^2)*1)+
    ((max(0,(x[10]/122 - 1))^2)*1)+
    ((max(0,(1-(x[10]+x[16]+x[20])/122))^2)*1)+
    ((max(0,(1-(x[6]+x[11]+x[13]+x[15])/3127))^2)*1)+
    ((max(0,(1-(x[7]+x[8]+x[9]+x[12]+x[14]+x[17]+x[21]+x[22])/5398))^2)*1)+
    ((max(0,((x[18]+x[19]+x[20])/9780 - 1))^2)*1)+
    ((max(0,(1-(x[10]+x[16]+x[18]+x[19]+x[20])/9780))^2)*1)+
    ((max(0,((x[7]+x[8])/1432 - 1))^2)*1)+
    ((max(0,(1 - (x[7]+x[8]+x[6]+x[15]+x[17]+x[21]+x[22]+x[16]+x[20])/1432))^2)*1)+
    ((max(0,(1 - (x[6]+x[15]+x[17]+x[21]+x[22]+x[16])/7046))^2)*1)
  
}
Problem$lb <- c(10,10,10,10,10,10,10,10,0,10,10,10,10,10,10,10,10,0,10,10,10,10)
Problem$ub <- c(1200000,1000000,1000000,1000000,1000000,1000000,1000000,1000000,1000000,1000000,1000000,1000000, 1000000,1000000,1000000,1000000,1000000,1000000,1000000,1000000,1000000,1000000)

outputfcn <- function(it, gbest, fx, x){
  cat(sprintf("Iteration: %d", it), sprintf("Error: %f", fx), "\n", sep = "\t");
  # return a negative value for PSwarm to abort execution
  1.0;
}


# Define a list with the solver options
Options <- list(
  cognitial = 0.5, fweight = 0.4, iweight = 0.9,
  maxf = 2000000000, maxit = 2000000000, size = 42 * Problem$Variables, iprint = 10000, social = 0.5,
  tol= 1E-6, ddelta = 0.5, idelta = 2,
  outputfcn = outputfcn, vectorized=0)

# Load the solver
#dyn.load("pswarm_r.so")  # Linux
dyn.load("pswarm_r")    # Windows

##########################
# Call the solver
Result <- .Call("pswarm_r", Problem, Options, .GlobalEnv)

# Results presents the obtained solution, its objective function value
# and a zero return value on success
print(Result)
View(Result[["x"]])
#########################
#########################

report <- data.frame(matrix(ncol = 23, nrow = 0))
for (i in 1:200){  
  # Call the solver
  set.seed(rnorm(1)*1E3)
  Result <- .Call("pswarm_r", Problem, Options, .GlobalEnv)
  cat("\n\nError=", Result$f, "\n\n")
  # print(Result)
  
  report<-rbind(report, append(Result$x, Result$f, after=0))
}

write.table(report, file = "./DG_011821.csv", fileEncoding = "UTF-16", na = "", sep = ",", row.names = F,
            col.names = c("error", "x1","x2","x3","x4","x5","x6","x7","x8","x9","x10","x11","x12","x13","x14","x15","x16","x17","x18","x19","x20","x21","x22"))

rm(i, Options, Problem, outputfcn, Result)
```
