## SupplementaryData for "Quantification of neuron types in the rodent hippocampal formation by data mining and numerical optimization": EC.R

```
# Define a list with the problem definition
rm(Problem, outputfcn, Options, Result)
Problem <- list()
Problem$Variables <- 70
Problem$objf <- function(x) {
  #################################
  ((((x[138]+x[139]+x[140]+x[141]+x[142]+x[143]+x[144]+x[145]+x[146]+x[147]+x[148]+x[149]+x[150]+x[151]+x[152]+x[153]+x[154]+x[155]+x[156]+x[157]+x[158]+x[159]+x[160]+x[161]+x[162]+x[163]+x[164]+x[165]+x[166]+x[167]+x[168]+x[169]+x[170]+x[171]+x[172]+x[173]+x[174]+x[175]+x[176]+x[177]+x[178]+x[179]+x[180]+x[181]+x[182]+x[183]+x[184]+x[185]+x[186]+x[187]+x[188]+x[189]+x[190]+x[191]+x[192]+x[193]+x[194]+x[195]+x[196]+x[197]+x[198]+x[199])/975656 - 1)^2)*10) +
  ((((x[141]+x[145])/71837 - 1)^2)*10) +
  ((((x[140]+x[144]+x[153]+x[190]+x[193])/192683 - 1)^2)*10) +
  ((((x[152]+x[154]+x[157]+x[158]+x[162]+x[164]+x[170]+x[194]+x[198])/116662 - 1)^2)*10) +
  ((((x[169]+x[174]+x[180])/41141 - 1)^2)*10) +
  ((((x[168]+x[173]+x[179]+x[183])/85605 - 1)^2)*10) +
  ((((x[178]+x[188])/45477 - 1)^2)*10) +
  ((((x[139]+x[143])/29955 - 1)^2)*10) +
  ((((x[138]+x[142]+x[147]+x[148]+x[149]+x[151]+x[189]+x[191]+x[192])/59890 - 1)^2)*10) +
  ((((x[146]+x[150]+x[155]+x[156]+x[159]+x[160]+x[161]+x[163]+x[167]+x[195]+x[196]+x[197]+x[199])/26574 - 1)^2)*10) +
  ((((x[166]+x[172]+x[177])/14325 - 1)^2)*10) +
  ((((x[165]+x[171]+x[176]+x[181]+x[182]+x[184]+x[185]+x[187])/17866 - 1)^2)*10) +
  ((((x[175]+x[186])/15279 - 1)^2)*10) +
  ((((x[141]+x[145])/99707 - 1)^2)*10) +
  ((((x[140]+x[144]+x[153]+x[190]+x[193])/150384 - 1)^2)*10) +
  ((((x[152]+x[154]+x[157]+x[158]+x[162]+x[164]+x[170]+x[194]+x[198])/88434 - 1)^2)*10) +
  ((((x[169]+x[174]+x[180])/42928 - 1)^2)*10) +
  ((((x[168]+x[173]+x[179]+x[183])/103964 - 1)^2)*10) +
  ((((x[178]+x[188])/123353 - 1)^2)*10) +
  ((((x[139]+x[143])/52181 - 1)^2)*10) +
  ((((x[138]+x[142]+x[147]+x[148]+x[149]+x[151]+x[189]+x[191]+x[192])/65671 - 1)^2)*10) +
  ((((x[146]+x[150]+x[155]+x[156]+x[159]+x[160]+x[161]+x[163]+x[167]+x[195]+x[196]+x[197]+x[199])/45851 - 1)^2)*10) +
  ((((x[166]+x[172]+x[177])/8198 - 1)^2)*10) +
  ((((x[165]+x[171]+x[176]+x[181]+x[182]+x[184]+x[185]+x[187])/17587 - 1)^2)*10) +
  ((((x[175]+x[186])/40382 - 1)^2)*10) +
  #ero
  ((((x[141]+x[145])/98156 - 1)^2)*1) +
  ((((x[140]+x[144]+x[153])/280413 - 1)^2)*1) +
  ((((x[152]+x[154]+x[157]+x[158]+x[162]+x[164]+x[170])/160326 - 1)^2)*1) +
  ((((x[169]+x[174]+x[180])/59349 - 1)^2)*1) +
  ((((x[168]+x[173]+x[179]+x[183])/132537 - 1)^2)*1) +
  ((((x[178]+x[188])/154135 - 1)^2)*1) +
  ((((x[139]+x[143])/88529 - 1)^2)*1) +
  ((((x[138]+x[142]+x[147]+x[148]+x[149]+x[151])/129640 - 1)^2)*1) +
  ((((x[146]+x[150]+x[155]+x[156]+x[159]+x[160]+x[161]+x[163]+x[167])/107061 - 1)^2)*1) +
  ((((x[166]+x[172]+x[177])/7745 - 1)^2)*1) +
  ((((x[165]+x[171]+x[176]+x[181]+x[182]+x[184]+x[185]+x[187])/20687 - 1)^2)*1) +
  ((((x[175]+x[186])/34963 - 1)^2)*1) +
  ((((x[190]+x[193])/7121 - 1)^2)*1) +
  ((((x[194]+x[198])/4718 - 1)^2)*1) +
  ((((x[189]+x[191]+x[192])/15768 - 1)^2)*1) +
  ((((x[195]+x[196]+x[197]+x[199])/9766 - 1)^2)*1) +
  (((x[207]/6583 - 1)^2)*1) +
  (((x[206]/9542 - 1)^2)*1) +
  ((((x[200]+x[201]+x[202]+x[203]+x[204]+x[205])/26691 - 1)^2)*1) +
  #
  ((((22*(x[146]+x[147]))/(12*(x[150]+x[151])) - 1)^2)*1)+
  ((((18*x[149])/(23*x[151]) - 1)^2)*1) +
  #re-interpreted
  (((x[149]/(0.81*(x[138]+x[142]+x[147]+x[148]+x[149]+x[151])) - 1)^2)*1) +
  (((x[151]/(0.17*(x[138]+x[142]+x[147]+x[148]+x[149]+x[151])) - 1)^2)*1) +
  (((x[153]/(0.10*(x[140]+x[144]+x[153])) - 1)^2)*1) +
  #(((x[149]/(0.81*(x[138]+x[142]+x[147]+x[148]+x[149]+x[151]+x[189]+x[191]+x[192])) - 1)^2)*1) +
  #(((x[151]/(0.17*(x[138]+x[142]+x[147]+x[148]+x[149]+x[151]+x[189]+x[191]+x[192])) - 1)^2)*1) +
  #(((x[153]/(0.10*(x[140]+x[144]+x[153]+x[190]+x[193])) - 1)^2)*1) +
  (((((7*x[144])/(15*x[153])) - 1)^2)*1) +
  ((((7*(x[169]+x[170]+x[162]+x[157]))/(8*x[153]) - 1)^2)*1) +
  ((((x[142]+x[144])/(2*(x[194]+x[195])) - 1)^2)*1) +
  ((((x[194]+x[195])/(x[192]+x[193]) - 1)^2)*1) +
  ((((x[192]+x[193])/(x[156]+x[157]) - 1)^2)*1) +
  ((((3*x[184])/(4*(x[171]+x[172]+x[173]+x[174])) - 1)^2)*1) +
  ((((2*(x[171]+x[172]+x[173]+x[174]))/(3*(x[176]+x[177]+x[179]+x[180])) - 1)^2)*1) +
  ((((15*x[182])/(38*x[171]) - 1)^2)*1) +
  ((((9*x[171])/(15*x[187]) - 1)^2)*1) +
  ((((4*x[183])/(21*x[173]) - 1)^2)*1) +
  ((((9*x[196])/(7*x[191]) - 1)^2)*1) +
  ((((9*x[161])/(23*x[156]) - 1)^2)*1) +
  ((((5*x[156])/(9*x[197]) - 1)^2)*1) +
  ((((3*x[197])/(5*x[150]) - 1)^2)*1) +
  (((x[151]/x[144] - 1)^2)*1) +
  (((x[144]/(2*x[166]) - 1)^2)*1) +
  ((((2*x[166])/(x[171]+x[172]) - 1)^2)*1) +
  ((((27*x[139])/(8*x[151]) - 1)^2)*1) +
  #121420
  ((((26*x[151])/(27*x[149]) - 1)^2)*1) +
  #121420
  ((((11*x[149])/(26*x[142]) - 1)^2)*1) +
  #121420 new below
  #((((11*x[151])/(27*x[142]) - 1)^2)*1) +
  ((((5*x[142])/(11*x[148]) - 1)^2)*1) +
  ((((14*x[148])/(5*x[156]) - 1)^2)*1) +
  ((((35*x[156])/(14*x[159]) - 1)^2)*1) +
  ((((5*x[159])/(35*x[155]) - 1)^2)*1) +
  ((((2*x[155])/x[163] - 1)^2)*1) +
  ((((11*x[163])/(10*x[160]) - 1)^2)*1) +
  ((((6*x[160])/(11*x[171]) - 1)^2)*1) +
  ((((27*x[171])/(6*x[185]) - 1)^2)*1) +
  ((((21*x[185])/(27*x[182]) - 1)^2)*1) +
  ((((14*x[182])/(21*x[181]) - 1)^2)*1) +
  ((((11*x[181])/(14*x[186]) - 1)^2)*1) +
  ((((3*x[186])/(11*x[175]) - 1)^2)*1) +
  ((((20*x[153])/(2*(x[140]+x[144])) - 1)^2)*1) +
  ((((13*(x[140]+x[144]))/(20*x[162]) - 1)^2)*1) +
  ((((5*x[162])/(13*x[158]) - 1)^2)*1) +
  ((((3*x[158])/(5*x[154]) - 1)^2)*1) +
  ((((4*x[154])/(3*x[168]) - 1)^2)*1) +
  ((((16*x[168])/(4*x[179]) - 1)^2)*1) +
  ((((7*x[179])/(16*x[188]) - 1)^2)*1) +
  #121420 Jones
  (((x[149]/(0.69*(x[138]+x[142]+x[147]+x[148]+x[149]+x[151])) - 1)^2)*1) +
  #Kim
  ((max(0,(1 - x[207]/114))^2)*1) +
  ((max(0,(1 - ((x[193]+x[190])/1751)))^2)*1) +
  ((max(0,(1 - (x[194]/2702)))^2)*1) +
  ((max(0,(1 - ((x[205])/1208)))^2)*1) +
  ((max(0,(1 - ((x[204])/2785)))^2)*1) +
  ((max(0,(1 - ((x[203])/1983)))^2)*1) +
  ((max(0,(1 -  x[206]/393))^2)*1) +
  ((max(0,(1 - ((x[189]+x[191]+x[192])/2865)))^2)*1) +
  ((max(0,(1 - ((x[195]+x[196]+x[199])/2738)))^2)*1) +
  ((max(0,(1 - ((x[202])/323)))^2)*1) +
  ((max(0,(1 - ((x[201])/530)))^2)*1) +
  ((max(0,(1 - ((x[200])/484)))^2)*1) +
  ((max(0,(1 - ((x[207])/1388)))^2)*1) +
  ((max(0,(1 - ((x[190]+x[193])/9990)))^2)*1) +
  ((max(0,(1 - ((x[194]+x[198])/9729)))^2)*1) +
  ((max(0,(1 - ((x[205])/5496)))^2)*1) +
  ((max(0,(1 - ((x[204])/14588)))^2)*1) +
  ((max(0,(1 - ((x[203])/15202)))^2)*1) +
  ((max(0,(1 - ((x[206])/1932)))^2)*1) +
  ((max(0,(1 - ((x[189]+x[191]+x[192])/8497)))^2)*1) +
  ((max(0,(1 - ((x[195]+x[196]+x[197]+x[199])/8575)))^2)*1) +
  ((max(0,(1 - ((x[202])/1534)))^2)*1) +
  ((max(0,(1 - ((x[201])/3771)))^2)*1) +
  ((max(0,(1 - ((x[200])/4539)))^2)*1) +
  ((max(0,(1 - ((x[207])/660)))^2)*1) +
  ((max(0,(1 - ((x[190]+x[193])/5212)))^2)*1) +
  ((max(0,(1 - ((x[198])/2661)))^2)*1) +
  ((max(0,(1 - ((x[205])/854)))^2)*1) +
  ((max(0,(1 - ((x[204])/1790)))^2)*1) +
  ((max(0,(1 - ((x[203])/1783)))^2)*1) +
  ((max(0,(1 - ((x[206])/810)))^2)*1) +
  ((max(0,(1 - ((x[189]+x[191]+x[192])/2536)))^2)*1) +
  ((max(0,(1 - ((x[195]+x[196]+x[197]+x[199])/1532)))^2)*1) +
  ((max(0,(1 - ((x[202])/115)))^2)*1) +
  ((max(0,(1 - ((x[201])/297)))^2)*1) +
  ((max(0,(1 - ((x[200])/549)))^2)*1) +
  ((max(0,(1 - ((x[190]+x[193]+x[194]+x[203]+x[204]+x[205]+x[207])/4718)))^2)*1) +
  ((max(0,(1 - ((x[189]+x[191]+x[192]+x[195]+x[196]+x[199]+x[200]+x[201]+x[202]+x[206])/10180)))^2)*1) +
  ((max(0,(1 - ((x[190]+x[193]+x[194]+x[203]+x[204]+x[205]+x[207])/12902)))^2)*1) +
  ((max(0,(1 - ((x[189]+x[191]+x[192]+x[195]+x[196]+x[199]+x[200]+x[201]+x[202]+x[206])/6051)))^2)*1)
}

Problem$lb <- rep(10, times = 70)
Problem$ub <- rep(1000000, times = 70)

outputfcn <- function(it, gbest, fx, x){
  cat(sprintf("Iteration: %d", it), sprintf("Error: %f", fx), "\n", sep = "\t");
  # return a negative value for PSwarm to abort execution
  1.0;
}


# Define a list with the solver options
Options <- list(
  cognitial = 0.5, fweight = 0.4, iweight = 0.9,
  maxf = 2000000000, maxit = 2000000000, size = 71 * Problem$Variables, iprint = 10000, social = 0.5,
  tol= 1E-6, ddelta = 0.5, idelta = 2,
  outputfcn = outputfcn, vectorized=0)

# Load the solver
#dyn.load("pswarm_r.so")  # Linux
dyn.load("pswarm_r")    # Windows

##########################
# Call the solver
Result <- .Call("pswarm_r", Problem, Options, .GlobalEnv)

# Results presents the obtained solution, its objective function value
# and a zero return value on success
#print(Result)
View(Result[["x"]])
###########################

report <- data.frame(matrix(ncol = 71, nrow = 0))
for (i in 1:100){  
  # Call the solver
  set.seed(rnorm(1)*1E3)
  Result <- .Call("pswarm_r", Problem, Options, .GlobalEnv)
  cat("\n\nError=", Result$f, "\n\n")
  # print(Result)
  
  report<-rbind(report, append(Result$x, Result$f, after=0))
}

#View(report)

#############################
write.table(report, file = "./ec022121_2.csv", fileEncoding = "UTF-16", na = "", sep = ",", row.names = F,
            col.names = c("error", "x138",	"x139",	"x140",	"x141",	"x142",	"x143",	"x144",	"x145",	"x146",	"x147",	"x148",	"x149",	"x150",	"x151",	"x152",	"x153",	"x154",	"x155",	"x156",	"x157",	"x158",	"x159",	"x160",	"x161",	"x162",	"x163",	"x164",	"x165",	"x166",	"x167",	"x168",	"x169",	"x170",	"x171",	"x172",	"x173",	"x174",	"x175",	"x176",	"x177",	"x178",	"x179",	"x180",	"x181",	"x182",	"x183",	"x184",	"x185",	"x186",	"x187",	"x188",	"x189",	"x190",	"x191",	"x192",	"x193",	"x194",	"x195",	"x196",	"x197",	"x198",	"x199",	"x200",	"x201",	"x202",	"x203",	"x204",	"x205",	"x206",	"x207"))

rm(i, Options, Problem, outputfcn, Result)
```
