## SupplementaryData for "Quantification of neuron types in the rodent hippocampal formation by data mining and numerical optimization": Sub.R

```
# Define a list with the problem definition
rm(Problem, outputfcn, Options, Result)
Problem <- list()
Problem$Variables <- 5
Problem$objf <- function(x) {
  ##################################
  (((x[133]+x[134])/462349 - 1)^2) +
  ((x[135]/49340 - 1)^2) +
  ((x[136]/16506 - 1)^2) +
  ((x[137]/7102 - 1)^2) +
  (((x[136]/38586 - 1)^2)*10) +
  ((((x[133]+x[134]+x[135])/263409 - 1)^2)*10) +
  (((x[133]/(10*x[134])) - 1)^2) +
  ((((x[133]+x[134]+x[135]+x[136])/731707)^2)*10) +
  ((((x[133]+x[134]+x[135]+x[136])/472500)^2)*10) +
  ((((x[133]+x[134]+x[135]+x[136])/792683)^2)*10) +
  ((((x[133]+x[134]+x[135]+x[136])/761590)^2)*10) +
  ((((x[133]+x[134]+x[135])/194457 - 1 )^2)*10) +
  (((x[136]/45940 - 1)^2)*10) +
  (((x[137]/33117 - 1)^2)*10) +
  (max(0,(1 - x[136]/89))^2) +
  (max(0,(1 - x[137]/235))^2) +
  (max(0,(1 - x[135]/18820))^2) +
  (max(0,(1 - x[136]/787))^2) +
  (max(0,(1 - x[137]/1852))^2) +
  (max(0,(1 - x[135]/41565))^2) +
  (max(0,(1 - x[136]/1110))^2) +
  (max(0,(1 - x[137]/1969))^2) +
  (max(0,(1 - x[135]/6279))^2) +
  (max(0,(1 - (x[135]+x[136]+x[137])/5301))^2) +
  (max(0,(1 - (x[135]+x[136]+x[137])/1109))^2)
}

Problem$lb <- rep(10, times = 5)
Problem$ub <- rep(1000000, times = 5)

outputfcn <- function(it, gbest, fx, x){
  cat(sprintf("Iteration: %d", it), sprintf("Error: %f", fx), "\n", sep = "\t");
  # return a negative value for PSwarm to abort execution
  1.0;
}


# Define a list with the solver options
Options <- list(
  cognitial = 0.5, fweight = 0.4, iweight = 0.9,
  maxf = 2000000000, maxit = 2000000000, size = 6 * Problem$Variables, iprint = 10000, social = 0.5,
  tol= 1E-20, ddelta = 0.5, idelta = 2,
  outputfcn = outputfcn, vectorized=0)

# Load the solver
#dyn.load("pswarm_r.so")  # Linux
dyn.load("pswarm_r")    # Windows

##########################
# Call the solver
Result <- .Call("pswarm_r", Problem, Options, .GlobalEnv)

# Results presents the obtained solution, its objective function value
# and a zero return value on success
print(Result)
###########################
report <- data.frame(matrix(ncol = 6, nrow = 0))
for (i in 1:200){  
  # Call the solver
  set.seed(rnorm(1)*1E3)
  Result <- .Call("pswarm_r", Problem, Options, .GlobalEnv)
  cat("\n\nError=", Result$f, "\n\n")
  # print(Result)
  
  report<-rbind(report, append(Result$x, Result$f, after=0))
}

write.table(report, file = "./Sub_012221.csv", fileEncoding = "UTF-16", na = "", sep = ",", row.names = F,
            col.names = c("error", "x133","x134","x135","x136","x137"))

rm(i, Options, Problem, outputfcn, Result)
```
